# SigFormer: an Attention-Based Framework for Robust Single-Sample Mutational Signature Decomposition

**DOI:** 10.64898/2026.01.20.700228

**Authors:** Yang Zhang, Muchun Niu, Chenghang Zong

## Abstract

Somatic mutational signatures imprint the history of exogenous exposures and endogenous processes on the genome, offering critical insights into pathologic etiology and disease risk. However, accurate signature decomposition at the single-sample level is still challenging when mutation burden is low, sampling noise is high, and candidate catalogs are large and redundant. Here, we present SigFormer, a set-conditioned transformer framework designed to facilitate robust somatic mutation analysis without reliance on large cohorts. By leveraging a cross-attention mechanism between customized reference input and sample mutation profile, SigFormer improves exposure recovery and detection accuracy compared with likelihood-driven refitting (MuSiCal) with the largest performance gains in high-noise and overcomplete settings. On PCAWG genomes, SigFormer preserves major tissue-level structure while sensitively and accurately capturing cooccurrence of low-abundance signatures but without the need for tumor-type–specific gating. In low-burden normal-tissue datasets spanning clonal expansion and microdissection studies, SigFormer maintains the high accuracy and recovers stable tissue-dependent patterns of SBS1/SBS5/SBS40a, pointing to underlying tissue-specific mutagenic heterogeneity in normal tissues. Finally, SigFormer quantifies an explicit unattributable residual component when the catalogue is incomplete, preventing forced allocation into flexible flat signatures and providing a useful signal for downstream analyses.

## Introduction

Somatic mutational signatures provide a compact and mechanistically interpretable readout of DNA damage and repair processes, capturing both intrinsic cellular activities^1,2^ and exogenous exposures^3,4^. By decomposing a sample’s mutational spectrum into contributions from distinct characteristic compositions (i.e. mutational signatures), we can link genome-wide mutation patterns to etiologic factors, disease mechanisms, and potential therapeutic vulnerabilities^1,5^. This computation procedure for mutational signature decomposition is commonly framed as estimating nonnegative activities of a reference catalog for each sample^6^. However, the process faces practical challenges including correlated signatures arising from noticeable degree of redundancy, low mutation burdens, and heterogeneous signal-to-noise across mutation classes^7^. Recent algorithmic development have advanced both usability and accuracy, for example, the SigProfiler ecosystem^8^ has emphasized robust single-sample attribution and probabilistic assignment of signatures to individual mutations through dedicated refitting frameworks. In parallel, MuSiCal^9^ introduced an integrated toolkit that enhances signature discovery and assignment, particularly by resolving the ambiguities associated with “flat” signatures and refining signature catalogs through large-scale reanalysis.

These advances have also been accompanied by more systematic benchmarking that clarifies when and why different tools succeed or fail. A recent comprehensive benchmark^10^ of signature fitting tools showed that performance depends strongly on mutation burden: SigProfilerSingleSample performs best at low mutation counts, whereas SigProfilerAssignment/MuSiCal performs best at high mutation counts. The study also cautioned that *ad hoc* restriction of the reference set can degrade performance. Complementing these observations, a large benchmarking study^11^ highlighted a key challenge for signature attribution: mutation spectra can often be reconstructed well by numerous different signature combinations, despite constraints that explicitly favor sparse solutions.

Thus, careful attention to stability and confidence is often required. Many pipelines still implicitly depend on cohort-scale discovery or strong sparsity assumptions, leading to reduced sensitivity for rare signatures or low-burden samples or both together. On the other hand, in relatively homogeneous cohorts or tightly constrained experimental designs, investigators often need principled ways to incorporate prior knowledge, for example, excluding implausible signatures or tailoring the reference set to the study context, to reduce destabilizing inference.

To approach this problem from a different angle, here we introduce a transformer-based framework for mutational signature decomposition, SigFormer, that treats the reference catalog as a customizable input set and uses cross-attention to aggregate evidence from a sample’s mutational profile for each candidate signature. This set-conditioned design naturally supports single-sample inference while allowing transparent, study-specific control of candidate signatures (e.g., masking or augmentation). We first benchmarked SigFormer on simulated catalogs spanning a wide range of mutation burdens, noise levels, and reference-set complexities, demonstrating that SigFormer outperforms the state-of-art algorithms across most comparison settings. We then validated its performance stability on PCAWG and biologically structured samples derived from microdissection and clonal expansion studies.

## Results

### Set-conditioned cross-attention enables single-sample mutational signature decomposition

Signature attribution is typically posed as estimating nonnegative activities over a fixed basis, but in practice it is often destabilized by two coupled difficulties: (i) correlated and redundant signatures that admit multiple near-equivalent reconstructions, leading to fragmented low-weight attributions, and (ii) variable signal-to-noise, particularly in low-burden samples where weak processes become hard to distinguish from sampling noise. To address these two challenges, we took advantage of the transformer-based machine learning algorithm^12^ and developed SigFormer, an AI architecture designed for single-sample mutational signature deconvolution under customizable reference catalogs (**Figure 1A**). In brief, SigFormer treats the reference catalog as an input set and uses set-conditioned cross-attention to match a sample’s observed mutation spectrum to candidate processes.

**Figure 1.**
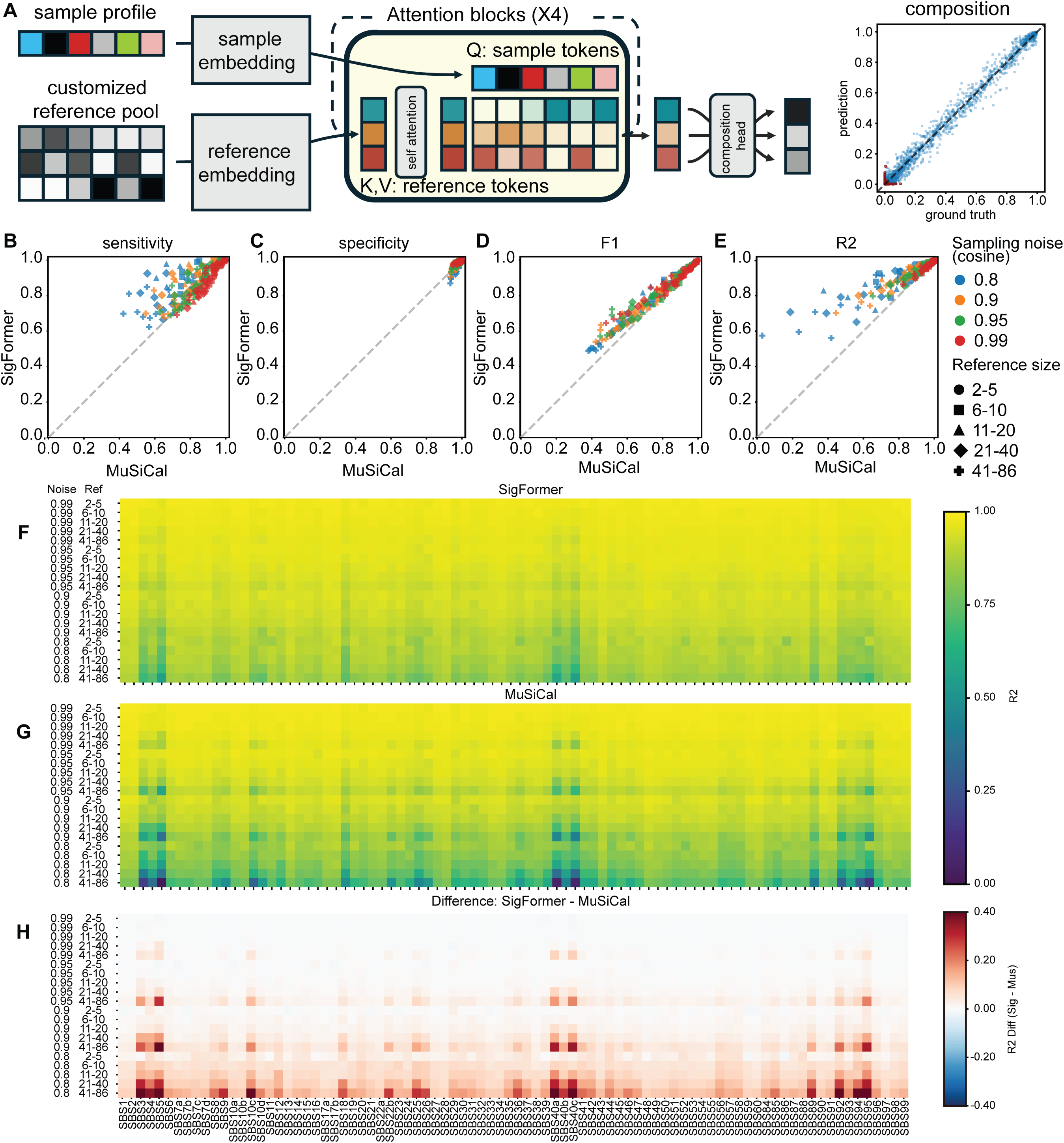
SigFormer architecture and performance on simulated dataset. (**A**) SigFormer architecture. Arrows indicate data flow. (**B-E**) General statistics across simulated datasets. Each dot represents a subset of 19,200 data, with color and marker type indicate data generation scheme. Sampling noise represents average sampling cosine similarity to probability matrix. Reference size indicates the number of reference signatures used to generate simulated data and used in the inference. Actual active signatures determined by target depth. (**F-H**) Spike-in test to assess performance across difference signature. SigFormer outperforms MuSiCal most significantly in flat signatures, for example, SBS3, SBS5, and SBS40a at higher noise scenario.

Specifically, SigFormer encodes each sample profile into a set of sample tokens, with each token representing an embedded mutation feature channel (for example, trinucleotide category or aggregated context features). Meanwhile, each candidate reference signature is encoded as one reference token, allowing the model to compare and attend to signatures at the token level. In each cross-attention block, sample tokens act as queries, while reference tokens provide keys and values, allowing each observed mutational context to retrieve a weighted combination of candidate signature representations that best match that context. Stacking multiple attention blocks enables iterative refinement: early layers form coarse matches that capture dominant processes, whereas later layers sharpen assignments through competition among similar signatures, effectively resolving ambiguity without relying on *ad hoc* catalog pruning. Because the reference set is explicitly represented as orderless tokens, SigFormer naturally supports variable-size and study-specific catalogs, thus allowing investigators to mask implausible signatures or augment the pool with custom candidates, all while maintaining a consistent inference procedure across samples.

To evaluate single-sample signature deconvolution under realistic sources of ambiguity, we constructed simulated profiles using a candidate reference pool comprising COSMIC v3.4 SBS signatures^13,14^ and an additional set of mock *de novo* signatures designed to span COSMIC-like feature statistics (e.g., entropy and sparsity, **Supplementary Methods**). For each condition, we sampled a reference set of variable size from small to large size of catalogs, generated activities under sparse-to-even compositions, and applied depth-controlled and Dirichlet-multinomial–controlled sampling noise to produce profiles with calibrated distortion from the noiseless expectation (**Supplementary Note 1** and **Supplementary Figure 1**).

Next, we averaged performance over 19,200 (3,000 batches with a batch size of 64) simulated samples per condition (**Supplementary Note 2**), with each point in **Figure 1B–E** representing a distinct combination of simulation settings (please see detailed breakdown in **Supplementary Figure 2**). Across conditions, performance was primarily governed by two factors: stochastic noise in the observed mutation profile and the candidate reference set size. This is consistent with known failure modes in signature attribution: higher noise increases sampling variance and reduces signal-to-noise, thus diminishing signature-defining features and lowering separability, which in turn decreases sensitivity for weak components. Furthermore, expanding the reference set worsens identifiability by adding correlated and potentially competing signatures, which further increases ambiguity and reduces attribution stability, even for dominant components. Consequently, performance degradation reflected a combination of reduced detectability under noise and increased identifiability limits from correlated or competing signatures (**Figure 1B**).

Overall, SigFormer showed greater robustness than a linear refitting baseline, preserving higher sensitivity over a broad range of noise and reference sizes. It is also worth noting that specificity remained high for both methods, with SigFormer showing a slight reduction relative to MuSiCal (**Figure 1C**). This pattern is consistent with a trade-off in which improved recovery of weak true positives can introduce minor “leakage” into correlated candidates when predictions are thresholded into discrete calls. With the result of the significantly improved sensitivity versus only marginal specificity loss, SigFormer achieved higher F1 scores in 157/180 conditions (**Figure 1D**).

Next, beyond binary detection, we assessed quantitative accuracy of inferred activities using the coefficient of determination (R²) between predicted and ground-truth compositions. R² provides a scale-free summary of how well the predicted exposure profiles explain the variance in the true activities, complementing detection metrics by emphasizing agreement in relative magnitude across signatures rather than their merely presence or absence. Overall, SigFormer achieved improved R² in 162/180 conditions (**Figure 1E**). The few settings in which SigFormer did not lead were confined to highly favorable, near-deterministic regimes with extremely low sampling noise and a small candidate signature set, the conditions under which strong prior constraints render the attribution problem particularly well-conditioned and thus allow linear mixture models to approach the optimal solution. Importantly, even in these cases, both SigFormer and MuSiCal operated near the performance ceiling (typically R² > 0.99), and the observed differences were minor (e.g., ΔR ² ≈ 0.01), suggesting practical convergence when the decomposition is unambiguous.

Next, to dissect quantitative accuracy at the level of individual signatures, we designed a spike-in benchmark. For each simulation setting, we generated simulated profiles by randomly sampling activities for the remaining n–1 active signatures (i.e., the background mixture varied across replicates) and then spiking in a target signature whose activity was titrated from 0% to 100% in 5% increments. To mimic small exposure fluctuations, the spike-in level for each replicate was perturbed with Gaussian noise (σ = 0.5%, truncated to [0, 100%]) before renormalization. We generated 100 repeats per condition to obtain robust performance estimates (**Supplementary Methods**). We then assessed performance using R² between predicted and ground-truth exposure vectors across titration levels for each signature and regime (**Figure 1F–H**).

Overall, both methods achieved high exposure concordance, with mean R² of 0.9228 for SigFormer and 0.8856 for MuSiCal. The dominant trends mirrored those observed in the global benchmark: profile noise and reference size were the primary drivers of performance across regimes. Despite these broadly shared regime effects, the identity of the underlying signatures also introduced substantial heterogeneity. In particular, flat and weakly distinctive signatures exhibited a marked reduction in exposure concordance (**Figure 1H**), consistent with increased ambiguity when there are competitions among correlated candidate pools. Signatures such as SBS3, SBS5, and SBS40a showed noticeably lower average R² (SigFormer 0.8495–0.8775; MuSiCal 0.7387–0.7992). Strikingly, these difficult signatures also corresponded to regimes where SigFormer most consistently exceeded MuSiCal, suggesting that set-conditioned attention enhances quantitative recovery precisely when signatures are most confusable within a fixed linear reference space.

Together, these simulated benchmarks demonstrate that single-sample signature deconvolution is dominated by two coupled challenges: reduced detectability under sampling noise and limited identifiability as the candidate pool expands with correlated competitors. Across a broad range of regimes, SigFormer consistently preserved higher sensitivity and F1 while preserving comparable specificity, indicating improved recovery of weak true components with only minor leakage into confusable candidates. Overall, SigFormer also achieved higher exposure concordance (R²) across most settings, with performance only converging in near-deterministic regimes where linear refitting approaches the ceiling.

### Benchmarking SigFormer on PCAWG genomes versus likelihood-driven MuSiCal assignments

After establishing SigFormer’s behavior on simulated benchmarks, we next asked whether a *prior-free*, single-sample inference model can produce biologically coherent mutational signature decompositions in real cancer genomes. This question is nontrivial, as many state-of-the-art pipelines implicitly or explicitly leverage prior expectations about which signatures should occur in which tumor types. As a reference, we compared SigFormer exposures to the MuSiCal reanalysis of the PCAWG cohort^15^. Here, we used the MuSiCal PCAWG signature assignments provided in Supplementary Table 4 from Jin *et al.*^9^ To minimize instability from shallow catalogs, we excluded ultra–low-burden samples (total SBS count <200).

Using reconstructed cosine similarity between observed 96-channel trinucleotide spectra and model reconstructions as a coarse goodness-of-fit measure, SigFormer achieved cosine similarity >0.9 across all retained samples and showed slightly higher cosine similarity than MuSiCal overall (**Figure 2A**). However, a higher cosine similarity does not necessarily imply a more faithful decomposition. In a highly overparameterized basis, improved reconstruction may instead arise from over-assignment of weak components to noise, a known failure mode in signature refitting, particularly for “flat” or correlated signatures. Therefore, we need to examine whether SigFormer’s exposure space preserves the structure of the raw mutational spectra rather than projecting samples into an overly discretized, prior-shaped signature manifold.

**Figure 2.**
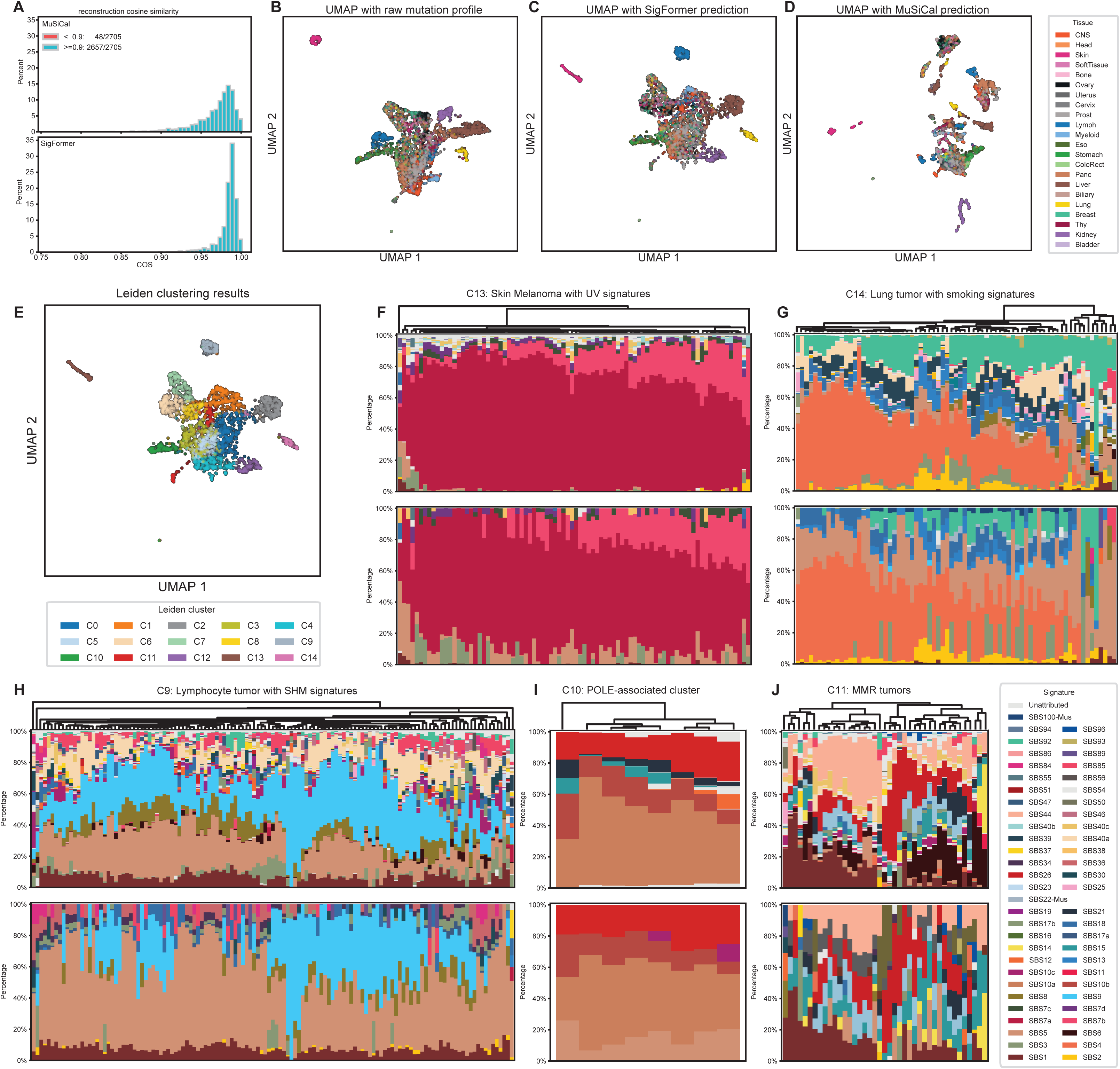
Benchmark of SigFormer on PCAWG data. (**A**) Reconstruction cosine similarity for MuSiCal (upper) and SigFormer (lower). (**B-E**) UMAP visualization of PCAWG samples. (**F-J**) Stacked barplot for predicted signature composition for Cluster: C9-11, C13, and C14. Upper panels are SigFormer results and the lower panels are MuSiCal result.

Here we performed UMAP visualizations using (i) raw 96-channel spectra, (ii) SigFormer exposures, and (iii) MuSiCal exposures in **Figure 2B-D**. SigFormer’s UMAP embedding largely recapitulated the geometry of the raw trinucleotide spectra, with major partitions corresponding to tumor primary tissue. This suggests that the inferred exposure representation remains anchored to the observed spectrum rather than drifting toward tissue-conditioned priors. By contrast, MuSiCal exposures produced a more discretized embedding, with sharper separation among subpopulations. This observation indicates that when signature refitting is performed on a curated basis, small changes in active signature sets can induce categorical shifts in exposure space.

Next, we dissected representative peripheral clusters (Leiden clusters as shown in **Figure 2E**), including ultraviolet-light (UV)-associated melanoma cluster (C13) (**Figure 2F**), smoking-enriched lung cluster (C14) (**Figure 2G**), lymphoid (CLL/BNHL) cluster (C9) (**Figure 2H**), POLE-driven cluster (C10) with residual SBS15/SBS21 (MMR-like) structure (**Figure 2I**), and heterogeneous MMR/MSI-like cluster (C11) across multiple adenocarcinoma types (**Figure 2J**). Meanwhile, other clusters dominated by broadly shared endogenous processes (C1-8 and C12) are detailed in **Supplementary Materials** (**Supplementary Notes 3** and **Supplementary Figure 3-13**).

### Benchmarking the performance in the UV-associated melanoma cluster

Cluster C13 in **Figure 2E** is a prominent cluster dominated by Skin-Melanoma samples and displays the expected UV signatures (**Figure 2F**). Both SigFormer and MuSiCal assigned high SBS7a and SBS7b, the dominant UV components in skin cancers^16^. Furthermore, SigFormer also recovered SBS38 in nearly all samples where MuSiCal assigned SBS38, consistent with prior reports that SBS38 is largely restricted to UV-associated melanomas and may reflect indirect UV-related damage.

Different from MuSiCal results, SigFormer reported consistent contribution of SBS7c and SBS7d (**Figure 3A**). Both have been proposed to arise from error-prone translesion synthesis opposite UV photodimers, making it plausible as a minor UV-associated component^17^. Because small components can sometimes reflect either genuine low-amplitude signal or flexible “noise absorption” among correlated signatures, to go beyond interpreting these differences solely from exposure frequencies, we performed a replacement-aware bootstrap necessity test to investigate whether each minor UV component can be substituted by its closest correlated candidates without degrading reconstruction evidence (**Supplementary Note 4**). Please refer to **Supplementary Methods** for the algorithmic details of replacement-aware bootstrap necessity test. For both SBS7c and SBS7d, the SigFormer predicted compositions positively correlate (spearman ρ =0.84 and 0.96, respectively, **Figure 3B**) showing that the bootstrap based necessity test support for retaining the signature. Samples with predicted composition >1% show both considerable gain of likelihood and significance, supporting that SBS7c and SBS7d indeed captures reproducible UV-consistent structure rather than acting as a generic noise sink.

**Figure 3:**
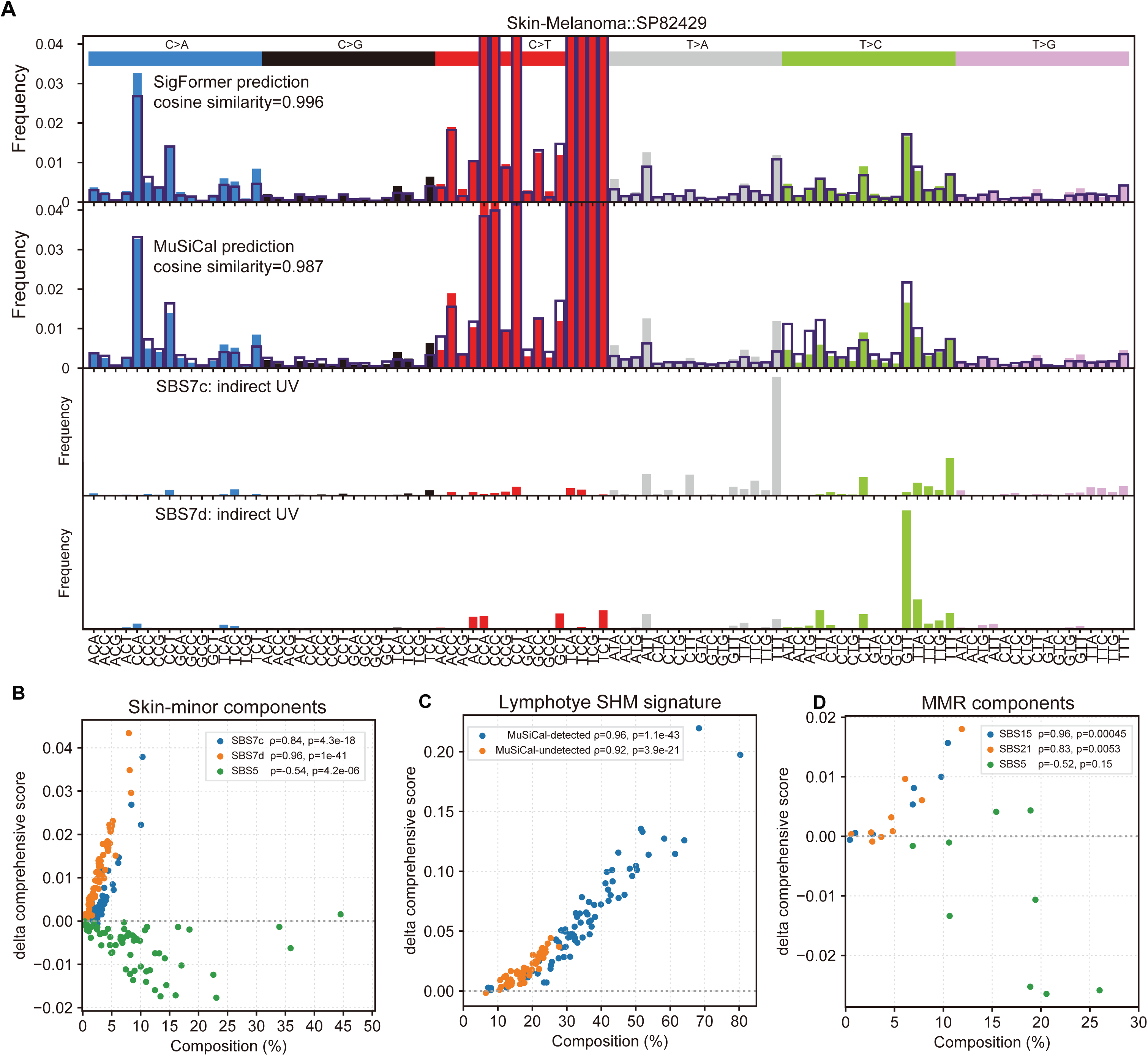
Evidence-based validation of low-abundance signature detection using a replacement-aware stability test **(A)** Mutation spectrum for a representative skin-melanoma sample. Colored bars show the observed (raw) mutation profile or COSMIC reference profiles, while outlines indicate reconstructions from the inferred signature compositions. From top to bottom: SigFormer reconstruction, MuSiCal reconstruction, and COSMIC profiles for SBS7c and SBS7d. (**B-D**) Replacement-aware stability test results validating low-abundance signature detection across the skin-melanoma cluster (C13), lymphocyte cluster (C9), and MMR-enriched cluster (C10).

Another notable divergence was the frequent MuSiCal assignment of SBS3/SBS5 within this melanoma cluster, whereas SigFormer more conservatively shifted comparable mass into an explicit “unattributed” component. When the basis is constrained and residual degrees of freedom are limited, SBS3 (homologous recombination deficiency)^18^ and SBS5 (clock-like, relatively flat)^2^ could be falsely over-assigned. In the same replacement-aware bootstrap necessity test, MuSiCal assigned SBS5 exhibited weak and often inconsistent evidence of necessity: the bootstrap-estimated per-mutation evidence change for retaining SBS5 was predominantly negative and showed only a moderate inverse association with the assigned SBS5 exposure (spearman ρ=−0.54, **Figure 3B**). This pattern is consistent with the disagreement being concentrated in the low-amplitude and broad-spectrum deviations where multiple flat signatures can trade off. In contrast, SigFormer can effectively reduce “garbage-bin” assignments into SBS5-like components when the evidence is lacking.

### Benchmarking the performance in the Smoking-enriched lung cluster

Cluster C14 in **Figure 2E** is dominated by Lung-SCC and Lung-AdenoCA and shows the expected smoking signature SBS4, consistently detected by SigFormer and largely concordant with MuSiCal. In **Figure 2G**, SigFormer additionally recovered SBS92 in multiple samples, a signature that has also recently been associated with tobacco smoking besides SBS4^3,19^, suggesting that tobacco mutagenesis may imprint more than one SBS component depending on tissue and exposure history.

Furthermore, SigFormer repeatedly favored SBS25 + SBS39 (**Figure 2G** upper panel), whereas MuSiCal instead assigned SBS22 together with its internally defined SBS100 (**Figure 2G** lower panel). This divergence is driven by two reproducible spectral deviations: (i) a prominent T>A peak at CTG, a feature for which the COSMIC catalogue provides essentially only SBS22 and SBS25 as plausible templates (**Supplementary Figure 15A**), and (ii) an enrichment of C>G substitutions, for which SBS39 is the only COSMIC signature with appreciable mass; MuSiCal compensates by introducing SBS100 to absorb this C>G-heavy component (**Supplementary Figure 15B**). Importantly, the mutually exclusive “explanations” provided by the two methods suggest that neither basis set may faithfully match the underlying process in these samples. Rather than interpreting SBS25/SBS22/SBS39/SBS100 labels etiologically, we view these assignments as a flag that a recurrent, poorly represented (or unknown) mutational process may exist in a subset of lung adenocarcinomas. This result shows that both SigFormer and MuSiCal could yield inconsistent and likely incorrect attributions, underscoring the limitations of reference-constrained decomposition. But on the other hand, the inconsistency indicates the lack of signature representing true generating process in the catalog.

### Benchmarking the performance in the Lymphoid (CLL/BNHL) cluster

Next, Cluster C9 in **Figure 2E** is enriched with Lymph-CLL and Lymph-BNHL samples and shows strong lymphocyte-origin processes. SigFormer consistently identified SBS9 (around 20–50% per sample, **Figure 2H**) and SBS9 is linked to polymerase-η–associated somatic hypermutation occurred in lymphoid cells^20^. In contrast, MuSiCal assigned SBS9 in only 50 out of 129 (39%) samples. To test whether SBS9 represents necessary, non-incidental structure rather than a flexible allocation choice, we applied the same evidence framework. In the replacement-aware bootstrap evidence test, retaining SBS9 yielded a positive improvement in reconstruction evidence (Δ comprehensive score) that increased monotonically with the SigFormer-inferred SBS9 activity. Importantly, this relationship was nearly indistinguishable between samples where MuSiCal reported SBS9 and those where it did not: both subsets showed strong positive rank correlations between SBS9 composition and Δ score (MuSiCal-detected: Spearman ρ = 0.96, p = 1.1×10⁻⁴³; MuSiCal-undetected: ρ = 0.92, p = 3.9×10⁻²¹) (**Figure 3C and Supplementary Figure 16**). The comparable evidence trends across the two groups suggest that SBS9-related signal is reproducible and not confined to MuSiCal-positive calls. Therefore, the MuSiCal “undetected” cases are more consistent with conservative calling or attribution ambiguity among correlated candidates, rather than a true absence of SBS9 in these lymphoid samples.

In addition to SBS9, SigFormer also assigned SBS85 more frequently than MuSiCal, which has been associated with AID-related mutagenesis in lymphoid cancers and often appears in clustered mutations. In MuSiCal, SBS9 and SBS85 appeared closer to mutually exclusive (either SBS9 *or* SBS85). While both signatures reflect related SHM/AID processes acting in the same cellular context, the exclusion indicates different potential processes. An alternative parsimonious explanation is identifiability: correlated lymphoid signatures can become substitutable under constrained refitting used in MuSiCal, yielding complementary assignments even when both processes contribute.

Furthermore, we observed increased SigFormer detection of SBS8 (hypothetically defective nucleotide excision repair)^21^ and SBS10c (defective POLD1 proofreading)^22^ in this cluster. Given that SBS10c is classically linked to polymerase proofreading deficiency rather than lymphoid SHM, this finding motivates a focused check for hypermutator phenotypes and strand-bias patterns in the underlying variant calls before attributing biology. In contrast, MuSiCal more frequently assigned SBS36 (MUTYH/BER deficiency)^23,24^, SBS17a/b (etiology uncertain; SBS17b has reported links to ROS damage and 5-FU in some contexts)^25^, and SBS34 (unknown). These components may represent genuine co-processes in subsets of samples, but they are also signatures prone to context-dependent misassignment when the spectrum is weakly informative, again emphasizing the need for targeted validation rather than relying on exposure prevalence alone for the relatively weak signals in signatures.

### Benchmarking the performance in the POLE-driven cluster with residual SBS15/SBS21 (MMR-like) structure

We observed two additional mismatch repair- (MMR)-related populations (C10 and C11 in **Figure 2E**). The first MMR-related cluster, which is dominated by the canonical POLE-associated trio SBS10a^26^, SBS10b^26^, and SBS28^27^, was comparatively “clean” (**Figure 2I**). In this cluster, MuSiCal consistently allocated ∼10–20% mass to SBS5, whereas SigFormer reassigned much of this residual component to SBS15 and SBS21, two MSI-associated signatures. To test whether these assignments reflect genuine residual structure rather than overfitting, we first examined the spectrum remaining after accounting for POLE signatures. After subtracting the reconstructed contributions of SBS10a/10b/28, the residual spectrum showed enrichment in C>T and T>C patterns and exhibited high cosine similarity to SBS15 (mean 0.90, range 0.82–0.96, **Supplementary Figure 17A**) and SBS21 (mean 0.82, range 0.64–0.93, **Supplementary Figure 17B**). We then performed necessity test comparisons. In the necessity test, excluding SBS15 or SBS21 from the basis produced a measurable loss of fit (Spearman ρ =0.96 and 0.83, respectively, **Figure 3D**), consistent with these components capturing reproducible residual structure. In contrast, SBS5 showed little evidence of necessity in the same analysis: despite non-trivial assigned mass by MuSiCal, the bootstrap-estimated per-mutation evidence change for retaining SBS5 was predominantly negative, with only two samples showing a clear positive gain (overall Spearman ρ = −0.52). Together, these results argue that the MSI-consistent components are unlikely to be incidental. This result supports the value of prior-free inference: unless there is explicit evidence to exclude a signature from the candidate basis, aggressive pre-pruning can predominantly redirect real, low-amplitude structure into flexible “catch-all” components (e.g., SBS5) rather than representing it as mechanistically informative signal.

### Benchmarking the performance in the Heterogeneous MMR/MSI-like cluster across multiple adenocarcinoma types

The second MMR-enriched cluster (C11 in **Figure 2E**) is comprised predominantly of adenocarcinomas arising from uterus, ovary, pancreas, colorectum, stomach, and biliary tract — cancer types in which mismatch-repair deficiency and microsatellite instability (MSI) are recurrently observed. Consistent with this etiology, SigFormer recovered a shared core set of MSI-associated signatures^28^ in almost all samples, most prominently SBS6, SBS20, SBS26, and SBS44, with additional low-to-moderate contributions by SBS14, SBS15, and SBS21 appearing in partially overlapping subsets (**Figure 2J**). Notably, SBS10c was detected only in a minority of cases and generally in a mutually exclusive manner relative to the strongest MSI components, suggesting sporadic polymerase-associated contributions superimposed on an MMR-deficient background. Despite the recurring MSI “core,” the overall decompositions in this cluster appeared heterogeneous, consistent with the well-known identifiability challenge among correlated MMR signatures (multiple MSI-associated bases can explain overlapping C>T / T>C structure with different fine-grained context preferences). In MuSiCal, the same sample group received more frequent allocations to SBS15, SBS21, and SBS26, while SBS44 was not reported. On the other hand, MuSiCal frequently assigned a signature labeled SBS97, reflecting catalog/version differences between MuSiCal’s customized reference set and the COSMIC definitions used by SigFormer. Direct comparison of spectra indicated that MuSiCal’s SBS97 is highly similar to COSMIC SBS44 (cosine similarity 0.85), and the similarity increased to 0.95 when restricting to C>T channels only, confirming that the two analyses are largely consistent but expressed under different signature nomenclature and minor spectral revisions.

### SigFormer yields stable and biologically coherent activities in low-burden normal datasets

To assess whether SigFormer remains robust in samples with low mutation burden, we next tested the performance of SigFormer for signature decomposition of the normal samples. Specifically, we assembled six published clonal expansion and microdissection datasets spanning multiple normal tissues, including blood, bone marrow^29^, breast^30^, colon^31^, intestine^32^, liver^33,34^, spleen and tonsil^29^. For each expanded clone or microdissected unit, we constructed a 96-channel mutation profile and inferred signature activities with SigFormer using a fixed reference catalogue. We then compared the intrinsic structure of samples in the original mutation-profile space with that in the inferred activity space, assessing whether activity-based representations preserve biologically expected relationships across tissues and sampling units.

UMAP embeddings constructed from raw 96-channel mutation profiles and from SigFormer-inferred activities yielded broadly concordant tissue-level organization (**Figure 4A and B**). In the raw 96-channel space, samples generally formed tight, well-separated tissue clusters, reflecting strong tissue-associated differences in the observed mutation spectrum (**Figure 4A**). Compared with the raw-profile embedding, the activity embedding provided a more interpretable and higher-resolution view of within-tissue heterogeneity, resolving multiple subclusters consistent with distinct mixtures of underlying mutational processes. Notably, SigFormer also revealed an additional cross-tissue continuum that was less apparent in the raw-profile representation, linking samples from multiple tissues along a shared axis of endogenous mutational activity.

**Figure 4.**
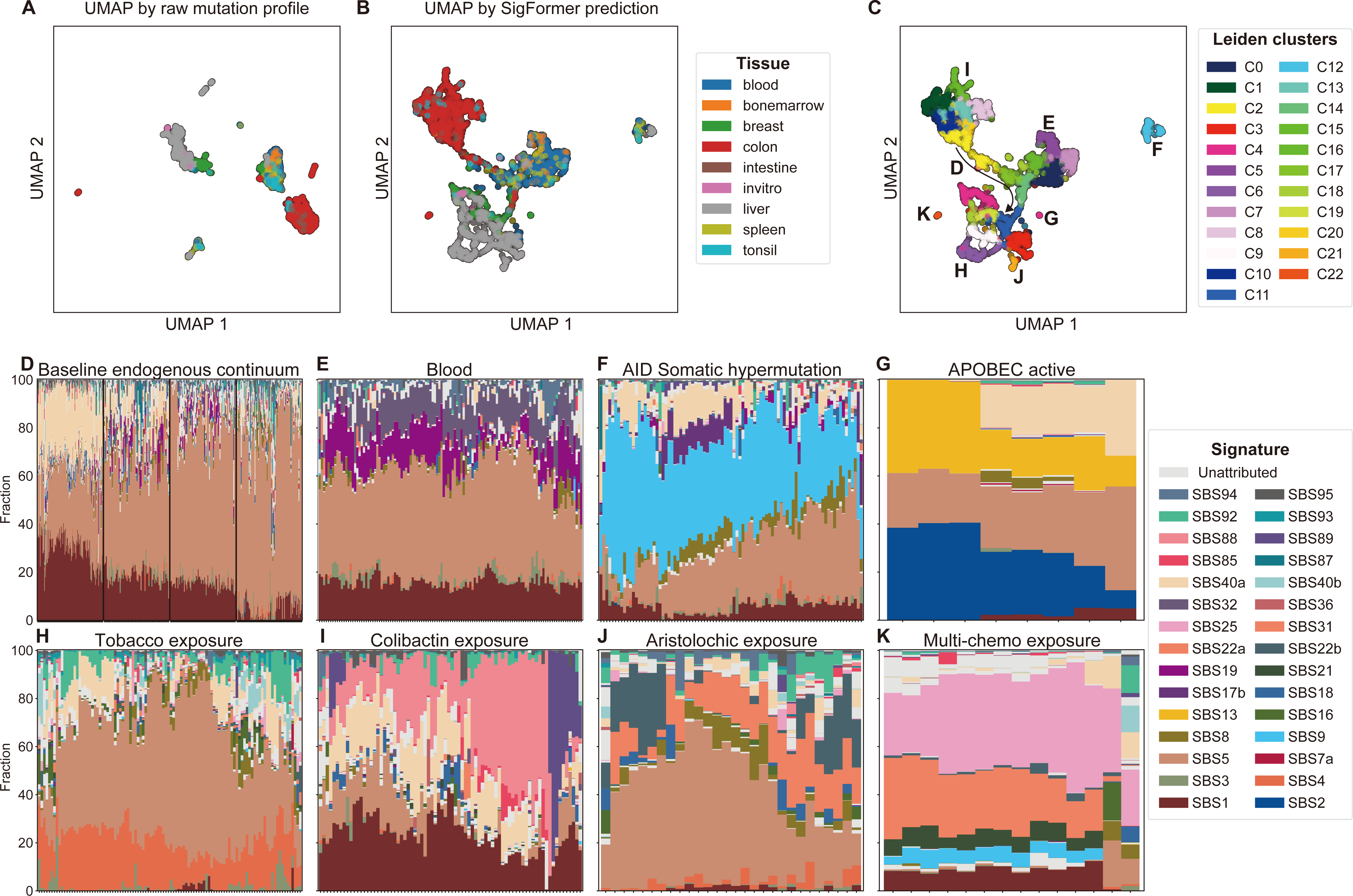
Performance of SigFormer in normal single cell expansion or microdissection samples. (**A-C**), UMAP dimension reduction showing that samples mainly cluster by tissue types. (**D-K**) SigFormer predicted signature compositions among different clusters.

To characterize this structure more systematically, we performed Leiden clustering on the activity embedding (**Figure 4C**) and summarized each community by its dominant signatures using stacked activity profiles (**Figure 4D–K**). The first major pattern corresponds to a “baseline endogenous continuum” spanning colon-, blood-, and liver-enriched regions, largely dominated by ubiquitous endogenous signatures with minimal contributions from overt exogenous or repair-deficiency processes (**Figure 4C and D**). Along this continuum, we observed a graded rebalancing among SBS1, SBS5, and SBS40a, that both distinguishes tissues and highlights shared baseline structure (**Figure 4D**). Colon-enriched clusters showed an approximately balanced mixture of SBS1, SBS5, and an SBS40-like component, consistent with high-turnover epithelia accumulating both CpG deamination-driven mutations (SBS1) and broadly clock-like background processes (SBS5/SBS40).

Progressing toward blood-enriched clusters, the relative contributions of SBS1/SBS40a-like activity decreased while SBS19 and SBS8 increased in a smaller cell portion, suggesting a shift toward hematopoietic-lineage–specific baseline damage. Notably, SBS19 has recently been linked to persistent endogenous lesions in hematopoietic stem cells, providing a plausible mechanistic basis for its selective enrichment in blood-derived samples. As spleen- and breast-derived samples became more represented along the same axis, SBS40a became nearly absent and activity redistributed across other signatures. Finally, liver-enriched clusters formed an endpoint with markedly reduced SBS1 (often <10%), consistent with substantial tissue-specific rebalancing of baseline processes, potentially reflecting differences in proliferative history, methylation/repair context, or the relative weighting of “flat” clock-like components.

Within blood and lymphoid lineages, SigFormer additionally recovered signatures aligned with known immune-specific mutagenesis. In B-lymphocyte-dominant communities (**Figure 4E**), SigFormer produced stable activity mixtures across many clones, echoing prior single-cell expansion studies of normal human lymphocytes^29^ while providing finer decomposition of broad SBSblood mutational patterns into multiple COSMIC components (SBS5, SBS19, SBS32, and SBS84 at 12:3:3:2 ratio). In memory B-cell–enriched clusters, SigFormer identified elevated SBS9 activity (**Figure 4F**), consistent with its established association with somatic hypermutation in lymphoid cells. We also observed an APOBEC-enriched community (**Figure 4G**), in line with reports^30^ of APOBEC-associated mutagenesis in breast-related clones and early evolutionary histories of breast cancer and related lineages.

Beyond endogenous variation, SigFormer was sensitive to exogenous exposures and tissue-specific mutagens. In donors with documented tobacco exposure, SigFormer detected co-occurring SBS4 and SBS92 activities, both of which are associated with tobacco-related mutagenesis (**Figure 4H**). In colorectal-derived clones, SigFormer recapitulated colibactin-associated mutagenesis through SBS88 and the SBS89 with unknown etiology as reported in normal intestinal crypts and colorectal epithelium by Henry et al^31^ (**Figure 4I**). In individuals with aristolochic acid exposure, SigFormer recovered SBS22a/b activity consistent with COSMIC annotations (**Figure 4J**).

Interestingly, SBS22a was also stably assigned in a lymphoma donor (**Figure 4K**) that had received combination chemotherapy, together with SBS9, SBS21, and SBS25. Although this mixture appears qualitatively plausible under a fixed reference catalog, closer inspection revealed a reproducible residual pattern, including a systematically underrepresented T>A signal at CTT, GTA, GTG, and GTT contexts (**Supplementary Figure 18 upper panel**). This observation shows the structure is not well captured by the available reference set. This is consistent with a common failure mode under catalogue incompleteness. The out-of-catalog processes are then likely approximated by a correlated combination of available signatures, yielding stable yet imperfect attributions. In the original study, Lee-Six et al. used hierarchical Dirichlet process (HDP) discovery to identify a de novo signature, SBSD, whose profile closely resembles a composite of SBS9, SBS21, SBS22a, and SBS25 (cosine similarity = 0.928, **Supplementary figure 18 lower panel**). Here when we replace these four catalog signatures with SBSD in the reference set, an improved reconstruction of the lymphoma spectra (mean cosine similarity improved from 0.943 to 0.978) was produced, supporting the interpretation that the apparent co-occurrence most likely reflects an out-of-catalog process rather than four independent activities.

Together, these analyses demonstrate that SigFormer produces stable and biologically coherent signature activity profiles across both clonal expansion and microdissection datasets. Across donors and tissues, it consistently recovers expected endogenous programs (including immune-associated processes) and sensitively captures established exogenous exposures, yielding interpretable tissue- and lineage-associated structures that parsimoniously explain the observed mutational spectra. In the vast majority of samples, residuals are minimal, supporting the adequacy of a reference-based decomposition. Only a small fraction of outlier communities, exemplified by the lymphoma donor above, show structured residual patterns suggestive of contributions not fully represented in the current reference panel. Thus, it is desired to incorporate an appropriate additional component into the reference pool to resolve these cases and restore high-fidelity reconstruction in future development of more comprehensive and adaptive reference frameworks.

## Discussion

Somatic mutational signature analysis is increasingly used as a mechanistically interpretable readout of DNA damage and repair, yet reliable single-sample decomposition remains difficult in the following regimes in practice: low mutation burden, heterogeneous noise, and inflated candidate catalogs containing correlated or weakly distinctive signatures. In this work, we introduced SigFormer, a set-conditioned transformer framework that treats the reference catalog as an explicit input set and uses cross-attention to aggregate evidence from an observed mutational spectrum into signature-specific activities. By construction, this design supports variable-size, study-specific candidate sets, including masking implausible signatures or augmenting the pool with custom candidates, while preserving a consistent inference procedure across samples.

SigFormer is designed around a central conceptual aim: prior-free single-sample inference. Here, “prior-free” does not mean the absence of assumptions altogether (the model is still conditioned on an input signature catalog), but rather the deliberate avoidance of tumor-type–specific gating, tissue-conditioned admissibility rules, or cohort-derived discovery steps that implicitly restrict which signatures are allowed to appear in a given sample. Instead, SigFormer performs inference by matching each sample’s observed 96-channel spectrum against a provided reference set, producing exposures that remain anchored to the data-generating spectrum rather than being discretized by curated, context-specific priors.

This property is particularly valuable for atypical cohorts, constrained experimental designs, and normal tissues, where strong tumor-type expectations may be unavailable, unreliable, or potentially misleading. Mechanistically, this outcome can be explained by the model’s set-conditioned attention architecture. Within the reference set, self-attention among signatures enables the network to represent redundancy and similarity relations, while the subsequent cross-attention from the sample to the reference set allocates explanatory mass under a normalized weighting scheme. In such a setup, highly similar (and therefore redundant) signatures naturally compete for the same evidence, causing their contributions to be mutually suppressive and sharpening the allocation toward the most consistent, data-supported decomposition rather than permitting diffuse mixtures across correlated candidates. Conversely, when the observed spectrum contains structure that is not well explained by any catalog element, SigFormer route probability mass into scattered mass that are collected as an explicit “unattributed” channel, effectively marking components as out-of-distribution relative to the provided reference set instead of forcing a correlated combination of imperfect proxies. In the PCAWG comparison, we have demonstrated that SigFormer maintained high reconstruction cosine similarity while producing an exposure representation that better preserved the geometry of the raw 96-channel spectra, consistent with exposures remaining spectrum-anchored rather than collapsing into a prior-induced manifold.

Beyond point estimates, we aimed to address a persistent interpretability failure mode in signature analysis: low-amplitude components can reflect genuine weak processes, but they can also arise as flexible allocations among correlated signatures that absorb noise. To avoid overinterpreting exposure frequencies, we introduced likelihood-based necessity and uniqueness tests that quantify per-sample evidence for individual signatures using refitted simplex-constrained exposures and parametric bootstrap. Necessity tests ask whether removing a target signature produces a reproducible loss of fit, while uniqueness tests ask whether the target provides a better explanation than any close alternative, explicitly correcting for the selection bias introduced by searching over replacements. Overall, this evidence layer is intended as a complement to neural predictions: it helps distinguish reproducible minor components from “noise sinks,” and it provides a principled way to support mechanistic claims in case studies without relying on exposure prevalence alone.

SigFormer also emphasizes explicit handling of catalogue incompleteness. In several real datasets, we observed structured residual patterns consistent with out-of-reference processes: when a spectrum cannot be faithfully represented by available signatures, stable but imperfect refits may instead distribute mass across correlated catalog components. By treating unexplained structure as informative rather than forcing attribution, a residual-guided workflow can flag candidate novel processes for targeted de novo extraction and cross-cohort validation. This is particularly relevant in normal tissues and treatment-exposed settings, where mutagenic processes could differ from those represented in cancer-dominant catalogs.

Several limitations should be noted. First, identifiability constraints are intrinsic to signature decomposition: highly correlated signatures can remain interchangeable under realistic noise, and no model can fully eliminate this ambiguity without additional information. SigFormer’s improved sensitivity in difficult regimes may therefore come with modest increases in “leakage” into correlated candidates when discrete presence/absence thresholds are applied.

Second, SigFormer is trained on simulated mixtures built from COSMIC signatures and synthetic feature-matched mock signatures, and its generalization depends on how well these simulations capture real-world artifacts and covariate structure (e.g., sequencing biases, context-dependent calling errors, and biological departures from simple multinomial sampling). Third, while the replacement-aware bootstrap necessity test provides principled support for minor components, they introduce additional computation and rely on modeling assumptions that may be imperfect in low-count regimes or overdispersed settings; extending these tests to richer count models (e.g., Dirichlet–multinomial likelihoods) is a natural direction.

Looking forward, several extensions could further strengthen single-sample mutational signature analysis. On the modeling side, integrating explicit count-likelihood objectives during training, improving uncertainty calibration of signature-level confidence, and extending the framework beyond 96-channel SBS spectra (e.g., incorporating indels, DBS, transcriptional strand, and genomic covariates) could broaden applicability. On the discovery side, systematic benchmarks under controlled catalogue incompleteness, coupled with residual-guided de novo extraction and reproducibility testing, could establish an end-to-end workflow for robustly identifying novel signatures while avoiding overfitting. Finally, connecting stable exposures to downstream likelihood models of gene- and region-level mutational risk in future would enable more direct integration of signature analysis with functional interpretation and disease mechanism studies.

## Acknowledgments

We are grateful to the McNair family for the support through McNair Scholarship. We thank other Zong lab members for their help in this project.

## Funding

C.Z. is supported by McNair Scholarship and SMaHT Program (1UG3NS132132 & 1UM1DA058229-01). Y.Z. is supported by pilot research funding from SMaHT Program.

## Author contributions

C.Z. and Y.Z. designed the project. Y.Z. performed the development and analysis. M.N. contributed to benchmarking through discussion. Y.Z. and C.Z. wrote the manuscript. C.Z. supervised the project.

## Competing interests

C.Z. is co-founders and equity holders of Pioneer Genomics Inc. The other authors declare no competing interests.

## Data and materials availability

Training data can be reconstructed by following the instructions provided in the project GitHub repository. PCAWG mutational data were downloaded from the ICGC Data Portal. Normal tissue datasets were obtained from the corresponding publicly available resources described in their respective publications.

## Code availability

The code for SigFormer training, simulation data generation, replacement-aware bootstrap necessity test, and other script used to generate figures for the paper has been deposited in the GitHub repository https://github.com/Yang-Zhang-717/SigFormer. The repository is currently private and will be made public before the publication of the manuscript.

## Supplementary Materials

### Methods

Supplementary Notes 1 to 4

Supplementary Figs. 1 to 18

