## Supplemental Materials for "SigFormer: an Attention-Based Framework for Robust Single-Sample Mutational Signature Decomposition"

**This file includes:**

**Methods**
**Supplementary Notes 1 to 4**
**Supplementary Figures 1 to 18**

### Methods

#### 1. Model architecture

SigFormerCore takes as input a sample mutational profile $x \in \mathbb{R}^{96}$ and a reference signature set $S \in \mathbb{R}^{N_{\text{ref}} \times 96}$. To stabilize scale across count and normalized inputs, the model computes depth $d = \sum_c x_c$ and forms a depth-normalized profile $x_{\text{nom}} = x/d \times 50{,}000$. Each of the 96 channels is treated as one sample token: a scalar "value embedding" (linear $1 \rightarrow d_{\text{model}}$) is summed with a learned channel embedding, yielding $T_{\text{smp}} \in \mathbb{R}^{96 \times d_{\text{model}}}$. Sample tokens are processed by an $n_{L,\text{smp}}$-layer Transformer self-attention tower (multi-head attention + MLP with residual connections and dropout). Each reference signature is projected by a linear layer $96 \rightarrow d_{\text{model}}$ to form reference tokens $T_{\text{ref}} \in \mathbb{R}^{N_{\text{ref}} \times d_{\text{model}}}$, which are updated by $n_{L,\text{ref}}$ stacked blocks consisting of (i) self-attention among reference tokens and (ii) cross-attention where reference tokens query the sample tokens (keys/values), producing sample-conditioned reference representations.

A composition head (LayerNorm + linear $d_{\text{model}} \rightarrow 1$) outputs per-signature logits mapped to a simplex using either softmax, entmax ($\alpha = 1.5$), sparsemax ($\alpha = 2$), or a learned entmax $\alpha$ clamped to $[1, 2]$. A confidence head is explicitly gradient-decoupled: it detaches the reference tokens and concatenates two global features, $\log(d)$ and a residual score $1 - \cos(x_{\text{nom}}, \hat{x})$ where $\hat{x}$ is the composition-weighted reconstruction, then predicts per-signature confidence via LayerNorm + linear + sigmoid.

#### 2. Training

Training uses on-the-fly simulation. A global reference bank contains COSMIC signatures plus synthetic de novo signatures generated with a cosine-similarity constraint (maximum similarity $\leqslant 0.6$) over a broad Dirichlet concentration range. For each batch, a subset of $K \in [10,90]$ reference signatures is sampled; in most batches a minimum COSMIC fraction is enforced to avoid degenerate de novo-only contexts. Synthetic samples are created by choosing a depth regime (low/medium/high) that caps the number of active signatures (4/10/16), sampling mixture weights from a Dirichlet distribution (with curriculum-controlled $\alpha$ bins), generating a clean profile as a weighted sum over the selected signatures, then adding profile noise (curriculum-controlled Dirichlet concentration) and sampling counts at the chosen depth. With probability norm_frac (default 0.25), the model receives the normalized noisy profile; otherwise it receives counts.

Optimization uses AdamW (default $d_{\text{model}} = 256$, 8 heads, $n_{L,\text{smp}} = 2$, $n_{L,\text{ref}} = 4$, dropout 0.1; batch size 64; 200 epochs). The learning rate follows per-batch warmup (first 5 epochs), a hold phase (until epoch 60), then linear decay to 0.05× base LR; gradients are clipped. The backpropagated loss is $\lambda_{\text{comp}} \text{MSE}(\widehat{w}, w) \cdot K + \lambda_{\text{conf}} \text{MSE}(\hat{c}, \exp(-s \mid \widehat{w} - w \mid))$. Reconstruction loss (depth-normalized MSE between $\hat{x}$ and noisy profile) is computed for monitoring but excluded from gradients in the provided script. Entmax $\alpha$ is not learned (LR=0), and the simplex is set to entmax15 from epoch $\geqslant 30$.

### 3. Simulated dataset

**Reference signature bank.**

We constructed a combined reference bank consisting of COSMIC v3.4 SBS signatures and 500 feature-matched mock de novo signatures. COSMIC signatures were taken as 96-channel probability vectors (trinucleotide contexts) and normalized to sum to one. Mock de novo signatures were generated to match the empirical distribution of COSMIC signature shape features (e.g., entropy and sparsity-related statistics), producing additional candidate processes that increase reference redundancy while remaining within a realistic morphological range.

**Factorial simulation design.**

We evaluated refitting performance across a factorial grid of four factors (Supplementary Figure 1): (i) **Reference pool size**, sampled from five bins: R2–5, R6–10, R11–20, R21–40, and R41–86; (ii) **Mixture concentration**, using Dirichlet concentration parameters $\alpha_{mix} \in \{0.1,1,10\}$ to generate sparse, intermediate, and near-uniform exposure regimes; (iii) **Mutation burden (depth)**, sampled from three ranges: 100–300, 1,000–3,000, and 10,000–30,000; (iv) **Profile perturbation (noise)**, modeled using a Dirichlet–multinomial process with concentration $\alpha_{noise} \in \{40, 120, 280, 2300\}$. This design yields 5 × 3 × 3 × 4 = 180 conditions.

**Reference pool construction per sample**

For each simulated sample, we first sampled a reference pool size K from the target bin and then selected K candidate signatures from the combined bank. Candidate sets were treated as the inference-time reference pool for both methods, ensuring a matched comparison under identical catalog complexity.

**Exposure sampling and ground-truth profile generation.**

Given a reference pool of K signatures, ground-truth exposures were drawn from a Dirichlet distribution with concentration α_mix (scaled by an all-ones vector of length K) and normalized to sum to one. The noiseless expected mutation profile was computed as the weighted mixture of reference signatures. A total mutation count (depth) was sampled uniformly from the target depth range.

**Dirichlet–multinomial sampling of observed profiles.**

To model over-dispersion and context-dependent sampling noise, we generated observed mutation counts using a Dirichlet–multinomial process. Specifically, a perturbed probability vector was drawn as p ~ Dirichlet(α_noise, p_true), followed by counts ~ Multinomial(depth, p). Across α_noise levels, this procedure produces increasing similarity between perturbed and noiseless profiles, corresponding to mean cosine similarity values of approximately 0.8, 0.9, 0.95, and 0.99, respectively.

**Spike-in benchmark for per-signature quantitative accuracy.**

For each simulation regime with an active signature set $\mathcal{S}(|\mathcal{S}| = n)$, we evaluated signature-wise accuracy using a controlled spike-in design. For each target signature $t \in \mathcal{S}$, we defined titration levels $\Gamma = \{0, 0.05, \ldots, 1.00\}$. At each replicate $r$, we first sampled a background exposure vector $\mathbf{b}^{(r)} \in \Delta^n$ over $\mathcal{S} \setminus \{t\}$(with $b_t^{(r)} = 0$), so that the remaining $n-1$active signatures varied across replicates. For each $\gamma \in \Gamma$, we perturbed the spike-in level by $\tilde{\gamma} =$

$\text{clip}(\gamma + \varepsilon, 0,1)$, where $\varepsilon \sim \mathcal{N}(0, \sigma^2)$with $\sigma = 0.005$, and formed the ground-truth exposure vector

$$\mathbf{a}^{(r,\gamma)} = (1 - \tilde{\gamma})\mathbf{b}^{(r)} + \tilde{\gamma}\mathbf{e}_t,$$

followed by renormalization to $\sum_k a_k = 1$ . The expected spectrum was $\mathbf{p}^{(r,\gamma)} = \sum_{k \in \mathcal{S}} a_k^{(r,\gamma)} \mathbf{s}_k$, from which counts were generated under the regime-specific noise model. We ran each inference method to obtain $\hat{\mathbf{a}}^{(r,\gamma)}$and quantified accuracy using

$$R^2 = 1 - \frac{\sum_{r,\gamma} \| \hat{\mathbf{a}}^{(r,\gamma)} - \mathbf{a}^{(r,\gamma)} \|_2^2}{\sum_{r,\gamma} \| \mathbf{a}^{(r,\gamma)} - \bar{\mathbf{a}} \|_2^2},$$

with 100 repeats per condition.

### 4. Replacement-aware bootstrap necessity test

To evaluate whether a candidate mutational signature is necessary and uniquely supported in a given sample (rather than merely a convenient surrogate), we developed the replacement-aware bootstrap necessity test. The test operates on three inputs: the observed trinucleotide mutation counts per sample $\mathbf{x}$(rows = samples, columns = 3-mer channels), a fixed reference signature dictionary $\mathbf{R}$(rows = signatures, columns = channels; each row normalized to a probability vector), and an initial exposure estimate $\mathbf{y}$(rows = samples, columns = signatures; row-normalized).

For each sample $s$and target signature $k$, we restrict analysis to signatures with estimated activity $y_{sk} \geq \tau$(default $\tau = 0.05$). The baseline reconstructed profile is

$$\mathbf{q}_{\text{raw}} = \sum_j y_{sj}\, \mathbf{R}_j, \sum_i q_i = 1,$$

and is compared to the normalized sample profile $\mathbf{p} = \mathbf{x}/\sum_i x_i$. To avoid depth-dependent scaling, all primary scores are computed in normalized space. We quantify reconstruction quality using (i) cosine similarity $\cos(\mathbf{p}, \mathbf{q})$and (ii) multinomial log-likelihood per mutation $\ell(\mathbf{p}, \mathbf{q}) = \sum_i p_i \log q_i$. Mean squared error is recorded but excluded from the primary score due to redundancy with cosine. The composite score is:

$$S(\mathbf{q}) = w_{\cos}\cos(\mathbf{p}, \mathbf{q}) + w_\ell \ell(\mathbf{p}, \mathbf{q}),$$

with $w_{\cos} = w_\ell = 1$by default.

To test replaceability, we generate alternative explanations by replacing the target signature mass $y_{sk}$with mixtures of $m \in \{1,2,3\}$candidate signatures selected from the top similar references (cosine similarity $\geq \theta$, default $\theta = 0.85$). The total exposure is conserved by construction, while mixture weights $\mathbf{w}$ are optimized on the simplex to maximize $S(\mathbf{q}_{\text{alt}})$under nonnegativity and per-component activity constraints (each component's final exposure $\geq \tau$). The best-performing alternative is defined as $\mathbf{q}_{\text{best}}$.

Statistical support is assessed by multinomial bootstrap: for $B$replicates (default $B = 30$), we sample $\mathbf{x}^{(b)} \sim \text{Multinomial}(N_{\text{eff}}, \mathbf{p})$and re-evaluate the score difference $\Delta^{(b)} = S(\mathbf{q}_{\text{raw}}) - S(\mathbf{q}_{\text{best}})$ (optionally re-optimizing $\mathbf{w}$per replicate). The one-sided empirical probability $p_{\text{alt}\geq\text{raw}} = \Pr(\Delta \leq 0)$determines necessity: a signature is called necessary/unique if $\Delta >$

0and $p_{\text{alt}\geq\text{raw}} < \alpha$(default $\alpha = 0.05$); replaceable if $\Delta < 0$and $\Pr(\Delta \geq 0) < \alpha$; otherwise inconclusive. Early stopping heuristics terminate bootstrapping when non-significance is evident, reducing compute without altering decision criteria.

### Supplementary Notes

#### 1. Empirical Grounding in PCAWG: Real-Data Distribution Summary

To generate simulated mutation profiles that are realistically anchored to real data distribution, we first characterized several distributional features of PCAWG samples using (i) raw 96-channel trinucleotide counts and (ii) signature exposure assignments obtained from Supplementary Table 4 of Jin et al. We defined **depth** as the total mutation count per sample (the sum across the 96 channels) and summarized its variability with a histogram on a log-scaled x-axis (**Supplementary Figure 1A**), reflecting the heavy-tailed burden landscape observed across donors and cancer types. Depth is not merely a nuisance variable: at low depth, multinomial sampling noise dominates, weak processes become statistically indistinguishable, and exposure estimates become inherently unstable; at high depth, subtle contributions become resolvable but a large reference pool can also "explain away" residual fluctuations by redistributing mass among correlated signatures, increasing the risk of overfitting in assignment.

Next, we quantified **signature sparsity** as the number of "active" signatures per sample, defining a signature as active if its exposure exceeded 3% (unless otherwise specified). Because detectability depends on count depth, we examined the relationship between active signature counts and depth. In PCAWG dataset, we observed a modest positive association (Spearman $\rho = 0.331$; **Supplementary Figure 1B**), consistent with the expectation that deeper samples permit the detection of additional low-level processes and allow optimization-based assignment to allocate small but nonzero exposures across more degrees of freedom. Notably, under extremely low depth (e.g., <200 total mutations), we observed a reversal of this trend: the estimated number of active signatures can increase rather than decrease. This behavior is consistent with two non-exclusive explanations: (i) stochastic fluctuation at very low counts can spuriously push multiple signatures above a fixed activity threshold, and (ii) the data may lack sufficient resolution to support stable sparse solutions, causing assignment algorithms to diffuse exposure across several correlated signatures in an attempt to reduce reconstruction residuals. In this regime, sparsity estimates should therefore be interpreted cautiously as they may reflect noise-driven dispersion rather than true biological complexity. Therefore in the main analysis, samples with mutation count <200 were excluded.

Finally, we summarized **composition evenness** by estimating a Dirichlet concentration parameter ($\alpha$) from exposure vectors restricted to active signatures and renormalized to sum to one (**Supplementary Figure 1C**). Intuitively, small $\alpha$ corresponds to "winner-takes-most" compositions dominated by one or two processes, whereas larger $\alpha$ reflects more even mixtures. This single-parameter summary provides an interpretable descriptor of how "spiky" or "diffuse" real tumor exposure profiles are and directly motivates the $\alpha$ settings used in our simulation grid to stress-test identifiability and robustness across realistic PCAWG-like regimes.

In addition to depth, sparsity, and compositional evenness, a critical determinant of decomposition performance is the *effective stability* of reference signatures in real tumors, i.e., how closely each reference approximates the latent processes operating in vivo. Direct quantification of "signature stability in true data" is generally infeasible: many signatures have unknown etiology, several are not cleanly separable due to collinearity, and tumor-specific context effects may systematically distort the observed spectrum relative to any fixed reference.

We therefore use **whole-profile reconstruction cosine similarity** as an empirically tractable proxy for the *average representativeness* of a reference bank under a given assignment method. Under an approximate generative model $x \approx \sum_k w_k s_k$, high reconstruction cosine indicates that (i) the reference spectra span the observed profiles and (ii) the estimated exposures provide a plausible mixture. Importantly, this proxy is **not** a direct measurement of per-signature truth: optimization against an expansive or redundant reference pool can inflate reconstruction similarity even when attributions are non-identifiable (a form of overfitting in the exposure space), thereby masking ambiguity rather than resolving it.

In PCAWG, MuSiCal achieves high-fidelity reference-based reconstructions, with $\sim 95\%$of samples exhibiting cosine similarity $> 0.9$(mean $\sim 0.96$; Figure 2A). This observation serves primarily as an empirical characterization of how closely real tumor spectra can be approximated by mixtures of known reference signatures. Building on this empirical fidelity distribution, we subsequently translate reconstruction variability into a controlled noise model for simulation: to emulate both stochastic sampling fluctuation and systematic deviation from a mean spectrum, we generate mutation counts using a Dirichlet–multinomial mechanism with concentration $\alpha_{\text{noise}} \in \{40,120,280,2300\}$, calibrated to yield average cosine similarity to the mean profile of approximately $0.8$, $0.9$, $0.95$, and $0.99$, respectively (**Supplementary Figure 1D**).

### 2. Benchmark metrics (classification-style and continuous accuracy)

We evaluated model performance using complementary metrics that capture both (i) detection of active mutational processes and (ii) quantitative accuracy of exposure estimates. Because simulated datasets provide known ground-truth compositions, errors can be directly attributed to omission (false negatives, FN) versus commission (false positives, FP), while continuous agreement can be assessed beyond binary activity calls.

To assess activity detection, we binarized each signature per sample as active/inactive under a chosen exposure threshold (e.g., ≥1%). We then computed standard confusion-matrix quantities: true positives (TP), false positives (FP), true negatives (TN), and false negatives (FN). From these we reported sensitivity/recall = TP/(TP+FN), specificity = TN/(TN+FP), precision = TP/(TP+FP), and F1 = 2·precision·recall/(precision+recall).

Across most simulated conditions, SigFormer showed markedly higher sensitivity than MuSiCal, recovering a larger fraction of truly active signatures. This improvement was most pronounced under high noise and large inference-time reference pools, where low signal-to-noise ratio and correlated signatures exacerbate identifiability challenges. Notably, the sensitivity gain depended on the underlying composition regime. In highly sparse compositions (e.g., one dominant signature), both methods performed well because the active set is readily identifiable. In contrast, sensitivity improvements were strongest in more even or multi-signature mixtures, where multiple moderate exposures lie near the detection boundary, and small underestimation can push true components below threshold, increasing FN. Consistent with this pattern, SigFormer's predicted exposures remained closely aligned with ground truth across increasingly challenging reference sets (**Supplementary Figure 2A–E**), whereas MuSiCal more frequently under-called weak-to-moderate components (**Supplementary Figure 2F–J**), resulting in higher FN rates after thresholding.

When examining specificity, SigFormer occasionally produced more low-composition predictions for truly absent signatures, yielding a modest FP increase relative to MuSiCal. This difference is best explained by distinct sparsification behaviors rather than "Transformer-specific" effects per se. MuSiCal, as a constrained likelihood-based optimization framework, often exhibits harder sparsification, frequently driving near-zero exposures to exact zeros under noisy or redundant reference settings. This suppresses FP calls after binarization and can inflate specificity, especially under low activity thresholds. In contrast, SigFormer predicts exposures through a learned mapping trained for robust reconstruction and composition fidelity across diverse conditions. Under high noise and large reference pools, SigFormer can exhibit small but non-zero leakage into correlated signatures, producing a tail of near-threshold FP values rather than confidently incorrect actives. Therefore, SigFormer's lower specificity is largely driven by many small-magnitude FP exposures, not gross misassignment of dominant processes. These trends are summarized in **Supplementary Figure 2K–N**, where SigFormer maintains consistently high specificity across the parameter grid, with the largest differences emerging in the most confounded settings (large reference, high noise).

Because sensitivity and specificity capture complementary failure modes, we emphasize F1 score as an integrated activity-detection metric. F1 balances precision and recall, penalizing both FP and FN, and avoids over-optimizing one axis (e.g., high specificity) at the expense of missing real processes. In our simulations, SigFormer's slightly reduced specificity was outweighed by its substantially improved sensitivity, yielding higher F1 in most settings (**Supplementary Figure 2M**). This trade-off is meaningful for mutational signature analysis, as failing to detect a true process can remove a mechanistic interpretation entirely, whereas low-amplitude FP predictions can be filtered downstream using exposure thresholds or stability criteria.

Binary activity calls are informative but discard magnitude information above threshold. Because signature decomposition is fundamentally a composition estimation problem, we additionally evaluated continuous accuracy using $R^2$ between predicted and true exposures, computed per signature across samples (with optional macro- or micro-averaging). $R^2$ captures both variance explained and systematic bias, distinguishing "roughly correct activity" from numerically accurate exposure estimates, which is critical for downstream interpretation (e.g., 5% vs 25% implies different mechanistic weight). Across nearly all simulated regimes, SigFormer achieved higher $R^2$ than MuSiCal, consistent with tighter agreement in the scatter plots (**Supplementary Figure 2A–E vs 2F–J**) and the summary heatmaps (**Supplementary Figure 2N**). Only under near-ideal conditions (extremely low noise and a reference set containing almost exclusively the true generating signatures) did MuSiCal occasionally attain marginally higher $R^2$; however, absolute $R^2$ values remained ≥0.99, making these differences practically negligible.

### 3. Mutational-signature landscape across the broadly shared endogenous broadly shared endogenous clusters in PCAWG

To complement the main text, we further characterized the mutational-signature distributions across the major clusters occupying the central "continent" region of the pan-cancer manifold (**Figure 2E**). In this region, clusters tend to be driven by varying mixtures of ubiquitous, clock-like processes (e.g., SBS1/SBS5/SBS40) together with tissue- or etiology-associated signatures that emerge along distinct manifold directions. Importantly, several signatures frequently observed in the central continent (including SBS5, SBS40, SBS18, and SBS8) are relatively "featureless" or "flat" in their 96-channel profiles, which can increase attribution ambiguity and make the inferred exposures more sensitive to model assumptions or reference-bank composition.

**Cluster C1: a multi-tissue "clock-like" core with sparse tissue-associated signatures**

The orange protrusion cluster C1 (**Figure 2E**) exhibits a comparatively featureless signature spectrum and contains a mixture of tissue types including CNS, blood-derived malignancies, and liver. Across both SigFormer and MuSiCal outputs, C1 is predominantly composed of SBS5, accompanied by lower-level SBS1 and scattered minor contributions from additional tissue-associated signatures (**Supplementary Figure 3**). This composition is consistent with SBS5 and SBS1 being widely observed "clock-like" processes present in many cancer types as well as in normal somatic lineages, with SBS5 showing a broad age correlation across tissues. Within hematopoietic samples, we additionally noted occasional enrichment of SBS9-like components. Canonically, SBS9 is proposed to reflect error-prone polymerase-η activity during somatic hypermutation, and has been most strongly linked to lymphoid lineages (e.g., subsets of CLL and B-cell malignancies). Therefore, any SBS9 attribution outside classical lymphoid contexts should be interpreted cautiously, as it may reflect tumor misannotation, mixed-cell contributions, or fitting instability when mutation burdens are low.

We also observed low-level SBS8 in a subset of samples within C1. Although SBS8 was historically labeled as "unknown etiology" in reference catalogues, multiple studies have connected SBS8-like spectra to nucleotide excision repair (NER) deficiency, implying that modest SBS8 activity may plausibly arise from impaired repair capacity in diverse tissues. At the same time, SBS8 is frequently discussed as a comparatively flat/low-specificity signature, increasing the risk of unstable refitting in low-mutation contexts.

**Cluster C0: mixed-tissue cluster with elevated SBS40a**

Adjacent to C1, the mixed-tissue cluster C0 (**Supplementary Figure 4**) exhibits a notable increase of SBS40a (often reaching ~30–40% in SigFormer assignments), accompanied by a shift in dominant primary tissues toward pancreas, breast, prostate, and liver. COSMIC annotations describe SBS40 as a broadly distributed signature of unknown etiology, with correlations to patient age reported in certain cancer types, supporting its interpretation as a clock-like component related to endogenous accumulation processes. However, SBS40 is also explicitly reported to be highly similar to SBS5, and this similarity can render the precise partitioning between SBS5- and SBS40-like activity uncertain, particularly in samples with limited mutation counts.

Interestingly, MuSiCal (lower panel) assigned appreciable SBS18 contributions across many C0 samples, whereas SigFormer did not show a comparable SBS18 trend, and the MuSiCal reconstructions exhibited lower cosine similarity on average in this region. SBS18 has been mechanistically linked to oxidative DNA damage (reactive oxygen species; ROS) and is often discussed together with SBS36 in the context of base-excision repair perturbations (e.g., MUTYH-related processes). Because SBS18 can co-vary with clock-like processes and may be hard to disentangle from other low-specificity components, broad SBS18 assignment may sometimes reflect attribution degeneracy rather than a true pan-tissue oxidative exposure spike. This type of method-dependent behavior is consistent with broader benchmarking literature showing that refitting outcomes can differ substantially across tools, especially for flat or weakly identifiable signatures.

**Cluster C2: liver-dominant smoking-associated group with SBS4/SBS92**

To the right of C0, the relatively isolated cluster C2 is largely composed of liver samples with strong smoking-associated exposures. Both SigFormer and MuSiCal detected elevated SBS4 as well as SBS92 in this group, consistent with tobacco-related mutational processes (**Supplementary Figure 5**). Notably, COSMIC commentary emphasizes that while tobacco smoking causes multiple cancer types, SBS4 has historically been most robustly detected in lung and head-and-neck cancers and is not always evident in other smoking-associated tumor types, implying that tobacco-related spectra may be tissue-context dependent. In contrast, more recent analyses have reported SBS4 presence beyond these classical contexts (including detection in liver in some datasets), highlighting ongoing refinements in how smoking-driven damage manifests across tissues and signatures.

We further observed that SigFormer's smoking-associated outputs were more continuous across samples, whereas MuSiCal showed greater discretization. Beyond the smoking axis, both methods also demonstrated sensitivity to sparse exposures: for example, a leftmost sample in this region showed strong SBS22a-like assignment by both methods. SBS22 is widely recognized as an aristolochic acid exposure signature and has been reported in liver and urinary-tract–related cancers. This supports the interpretation that even within a cluster dominated by smoking-associated mutagenesis, rare but high-confidence environmental signatures can remain detectable.

**CNS-associated cluster C11 and the SBS8-enriched manifold extension toward C8**

The upper-left manifold branch includes the red cluster C11, dominated by CNS samples. Alongside the expected clock-like background (SBS1/SBS5/SBS40a), both SigFormer and MuSiCal detected an SBS8-enriched subset (**Supplementary Figure 6**). As described above, SBS8 has been repeatedly linked to NER deficiency, providing a plausible mechanistic interpretation for SBS8 enrichment when present in CNS tumors, although the aetiology remains heterogeneous and attribution stability can be challenging in low-mutation contexts.

Continuing along this manifold direction leads to C8 (**Figure 2E**), primarily composed of ovary and prostate tumors. In this region we observed (i) a higher SBS8 detection rate, and (ii) consistent SBS3 contributions in a subset of clusters (**Supplementary Figure 7**). SBS3 is strongly associated with homologous recombination deficiency (HRD), including germline/somatic BRCA1/BRCA2 alterations and BRCA1 promoter methylation, and is classically enriched in ovarian, breast, and pancreatic cancers. Thus, the presence of SBS3 within ovary-containing substructures is consistent with well-established HRD biology.

In ovary samples, we additionally observed increased C>G patterns localized to CpG-rich contexts, interpreted as SBS39-like in SigFormer and as a study-defined component ("SBS100") in MuSiCal. COSMIC currently lists SBS39 as a signature of unknown etiology, and it has not reached the same consensus interpretability as SBS3, SBS4, or SBS22. Accordingly, while the observed motif-level enrichment suggests a reproducible process within this manifold branch, the mechanistic labeling of such patterns should be treated as provisional until supported by orthogonal evidence (e.g., genomic topography, strand bias, or experimental replication), particularly given the known instability of certain low-specificity signatures during refitting.

**Distinct process-dominated clusters: HRD (C6) and APOBEC (C7)**

Two clusters stand out as process-dominated "islands" in the manifold. C6 is enriched for HRD-associated SBS3, consistent with COSMIC annotations and prior pan-cancer analyses linking SBS3 to BRCA-related HR repair failure (**Supplementary Figure 8**). Meanwhile, C7 is dominated by APOBEC-associated SBS2/SBS13 activity, which is widely attributed to AID/APOBEC cytidine deaminase mutagenesis and frequently co-occurs as paired signatures in the same samples (**Supplementary Figure 9**). Although APOBEC activation is common across cancers, its triggers are context-dependent (viral infection, inflammation, replication stress, or retrotransposition have been proposed), and COSMIC notes that definitive causal evidence remains incomplete for many settings.

**Central clusters C3 and C5: balanced clock-like mixtures with higher SBS18 prevalence in C3**

At the center of the continent, clusters C3 and C5 both show relatively balanced mixtures of SBS1, SBS5, and SBS40a (**Supplementary Figure 10 and 11)**. Within this balanced region, C3 exhibits a higher prevalence of SBS18 compared to the prostate/pancreas-enriched C5. SBS18 has been observed across diverse tumor types and is mechanistically connected to oxidative damage, supporting the hypothesis that SBS18-like activity may arise in multiple tissues rather than being strictly tissue-restricted. This multi-tissue presence is consistent with the interpretation that C3 represents a heterogeneous tissue mixture with shared endogenous/oxidative influences, whereas C5 reflects more lineage-constrained tumor composition with relatively stable clock-like baselines.

**Gastrointestinal extension: C10 enriched for SBS17a/b and SBS41, and a gastric SBS93 micro-cluster**

Extending outward from C3, cluster C10 is primarily composed of esophagus and stomach samples, with increased SBS17a and SBS17b contributions and detectable SBS41

(**Supplementary Figure 12**). Multiple studies have linked SBS17-family signatures to gastrointestinal contexts, including Barrett's esophagus and esophageal adenocarcinoma, with proposed mechanisms involving chronic inflammation, reflux of acid and bile, and persistent oxidative stress; additionally, SBS17b has been discussed in relation to therapy-associated processes in some settings. By contrast, SBS41 remains of unknown aetiology in COSMIC, and thus its interpretation is currently limited to descriptive reporting rather than mechanistic inference.

On the right side of this gastrointestinal region, we identified a smaller stomach adenocarcinoma cluster with consistent SBS93 assignment by both MuSiCal and SigFormer. COSMIC reports SBS93 as being almost exclusively identified in gastric cancers in PCAWG, with additional detection in esophageal squamous cell carcinomas, and de novo extraction efforts have similarly emphasized its strong stomach association. This agreement across methods and external datasets supports SBS93 as a robust gastric-associated process within the manifold.

**Kidney island: SBS40c as a renal-restricted component**

Finally, the far-right "island" is dominated by kidney samples, in which we observed SBS40c almost exclusively (**Supplementary Figure 13**). COSMIC specifically notes that SBS40 can be split into SBS40a/b/c based on large-scale clear cell renal cell carcinoma (ccRCC) analyses, and that SBS40c has been identified in ccRCC with improved specificity relative to the unsplit SBS40 reference. Moreover, SBS40b has been reported as a kidney-associated signature linked to epidemiologic variation in ccRCC incidence and to markers of reduced kidney function, supporting the broader concept that SBS40-family subcomponents can capture renal-specific mutagenic processes.

### 4. Evidence-based validation of low-abundance signature detection using a replacement-aware stability test

Across multiple tumor clusters, we observed strong agreement between SigFormer and MuSiCal in dominant mutational processes, yet systematic discrepancies emerged in low-to-moderate exposures. For instance, in Skin–Melanoma (**Figure 2F**), SigFormer consistently assigned SBS7c/SBS7d at modest levels (2–5%) across most tumors, whereas MuSiCal detected these UV-related components only sporadically and instead attributed 5–10% exposure to SBS5 in the majority of samples. Manual inspection of the underlying 96-channel spectra supported the SigFormer interpretation. In one representative example (**Figure 3A**), we observed enrichment of T>A substitutions consistent with SBS7c and a characteristic T>C pattern aligned with SBS7d. Critically, SigFormer more accurately matched these localized peaks than MuSiCal, suggesting that some MuSiCal SBS5 assignments may function as a "mass absorber" rather than reflecting a truly active etiology.

To rigorously assess whether such detections are statistically credible, we developed a replacement-aware evidence test for signature necessity and uniqueness. The central hypothesis is that a genuine mutational process should be difficult to replace without degrading the likelihood of the observed mutation accumulation, even when the total exposure mass is preserved. For each sample, we first reconstruct the spectrum from the reported signature composition and quantify reconstruction quality using three complementary metrics: (i) **Spectrum Cosine Similarity (SCS)** in normalized probability space, (ii) **Normalized Mean Squared Error (nMSE)**, and (iii) **Per-mutation Log-likelihood (pLL)** under a multinomial model. Since nMSE is typically highly redundant with SCS, we define a composite reconstruction score dominated by orthogonal evidence channels:

$$\mathrm{CRS} = w_{\mathrm{SCS}} \cdot \mathrm{SCS} + w_{\mathrm{pLL}} \cdot \mathrm{pLL},$$

with nMSE retained as a diagnostic output but not double-counted in the score. Importantly, pLL is normalized per mutation, making CRS comparable across samples of varying sequencing depth.

For each target signature deemed active (composition ≥5%), we identify its most similar candidate signatures (cosine similarity ≥0.85) and generate alternative explanations by replacing the target with a mixture of one to three candidates while keeping total exposure mass constant. Mixture weights are optimized on the simplex to maximize CRS, with an additional feasibility constraint that every component in a multi-signature alternative remains active in the final composition (≥5%), preventing degenerate solutions driven by infinitesimal weights.

We then quantify robustness via **Multinomial Bootstrap Replacement Test**. We generate bootstrap replicates by resampling mutations from the observed spectrum (multinomial sampling at effective depth) and recompute CRS for the original and best alternative models. The empirical p-value is estimated as the fraction of replicates where the alternative matches or outperforms the original, enabling a decision of **necessary/unique**, **replaceable**, or **inconclusive**. Early-stopping heuristics reduce compute when effects are clearly non-significant.

We validated replacement test using a label-shuffling control in Cluster 5 (**Supplementary Figure 11**), containing SBS1, SBS5, and SBS40a. Under correct labeling, all three signatures showed positive correlations between exposure and CRS (**Supplementary Figure 14**). As expected, the sparse SBS1 exhibited the strongest monotonicity (Spearman 0.90), followed by SBS40a (0.83). SBS5 performed worst, consistent with its flatter profile and higher redundancy (≥2 COSMIC signatures with cosine similarity >0.85). In contrast, shuffled assignments produced markedly lower, typically negative CRS, indicating that incorrect labels are systematically replaceable by better-fitting alternatives.

We then applied this framework to three clusters showing pronounced SigFormer–MuSiCal disagreement. In Skin-Melanoma, SigFormer-detected SBS7c and SBS7d exhibited significant positive exposure–CES correlation despite low abundance, consistent with real UV-derived processes (**Figure 3B**). Conversely, MuSiCal's SBS5 allocations often produced negative CES, supporting the interpretation of SBS5 as a mass absorber in these samples.

In lymphocytes, MuSiCal failed to detect active SBS9 in nearly half of samples, particularly at intermediate exposures (10–25%) (**Figure 2H**). Meanwhile, exposure–CES trends did not separate detected versus undetected groups (**Figure 3C**), and spectra displayed clear residual SBS9-like features (**Supplementary Figure 16**), supporting true SBS9 activity even when MuSiCal reports none.

Finally, in the isolated C10 sub-cluster (**Figure 2I**), SigFormer frequently reported co-occurrence of SBS15 and SBS21, both associated with mismatch repair deficiency (dMMR). Although full-spectrum cosine similarity was high for both methods (SigFormer 0.997, MuSiCal 0.992) (**Supplementary Figure 17A and B**), residual inspection after subtracting POLE-related components (e.g., SBS10a/b and SBS28) revealed clear SBS21-like structure and additional evidence for SBS15. Both signatures showed strong exposure–CES correlations (SBS15 $\rho=0.95$; SBS21 $\rho=0.83$) (**Figure 3D**), indicating that high global similarity can mask biologically meaningful substructure unless targeted necessity testing is applied.

Together, these results demonstrate that SigFormer's low-to-moderate abundance detections are frequently supported by spectral features and validated by a statistically grounded necessity-and-uniqueness test. Beyond this study, the framework provides a general tool to assess whether a reported signature is genuinely required by the data or replaceable under realistic sampling noise, enabling more reliable downstream etiological interpretation.

**Supplementary Figures:**

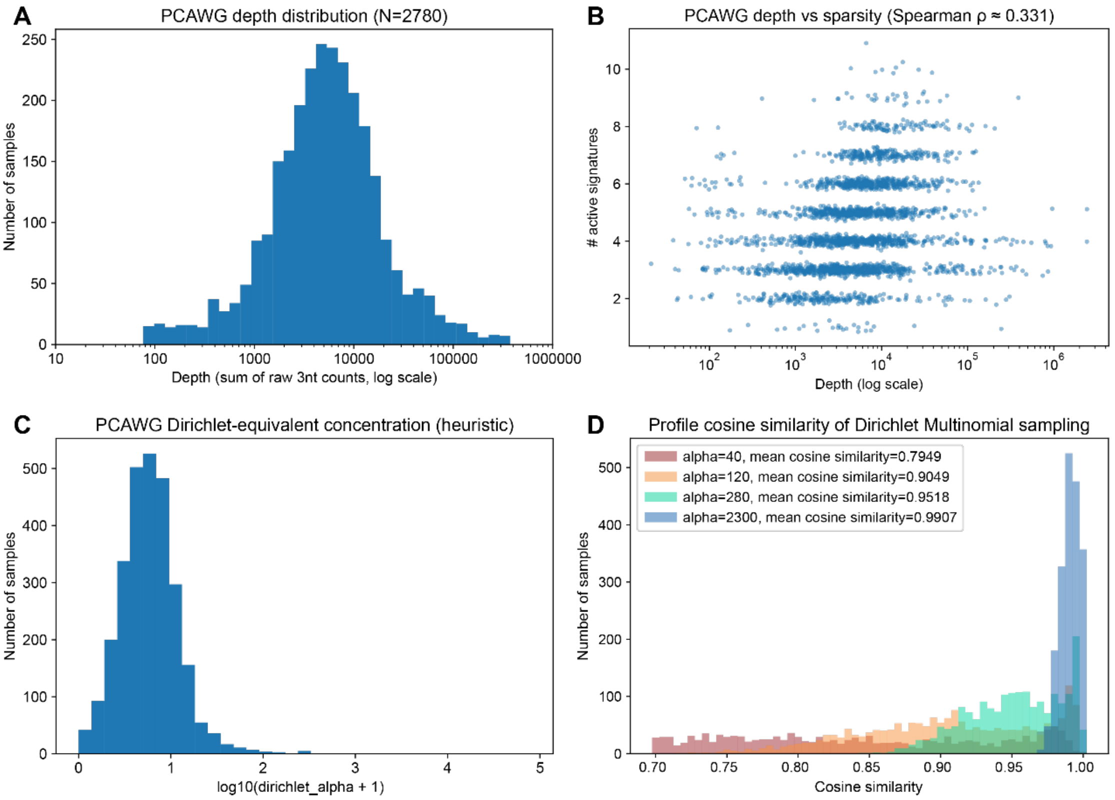


**Supplementary Figure 1: Dataset-level depth, sparsity, composition heterogeneity, and sampling robustness in PCAWG.**

(**A**) Distribution of mutational profile depth across PCAWG samples (N = 2,780), where depth is defined as the total number SNVs per sample (log scale). (**B**) Relationship between depth and signature sparsity, quantified as the number of active signatures per sample; depth and sparsity show a modest positive association (Spearman ρ = 0.331). (**C**) Effective Dirichlet concentration inferred from per sample composition, shown as log10(α + 1), where lower α indicates sparser mixtures and higher α indicates more diffuse compositions. (**D**) Cosine similarity distribution for samples generated under different Dirichlet concentration parameters (α) using Dirichlet–multinomial sampling. This distribution is used to calibrate the noise level when constructing simulated datasets

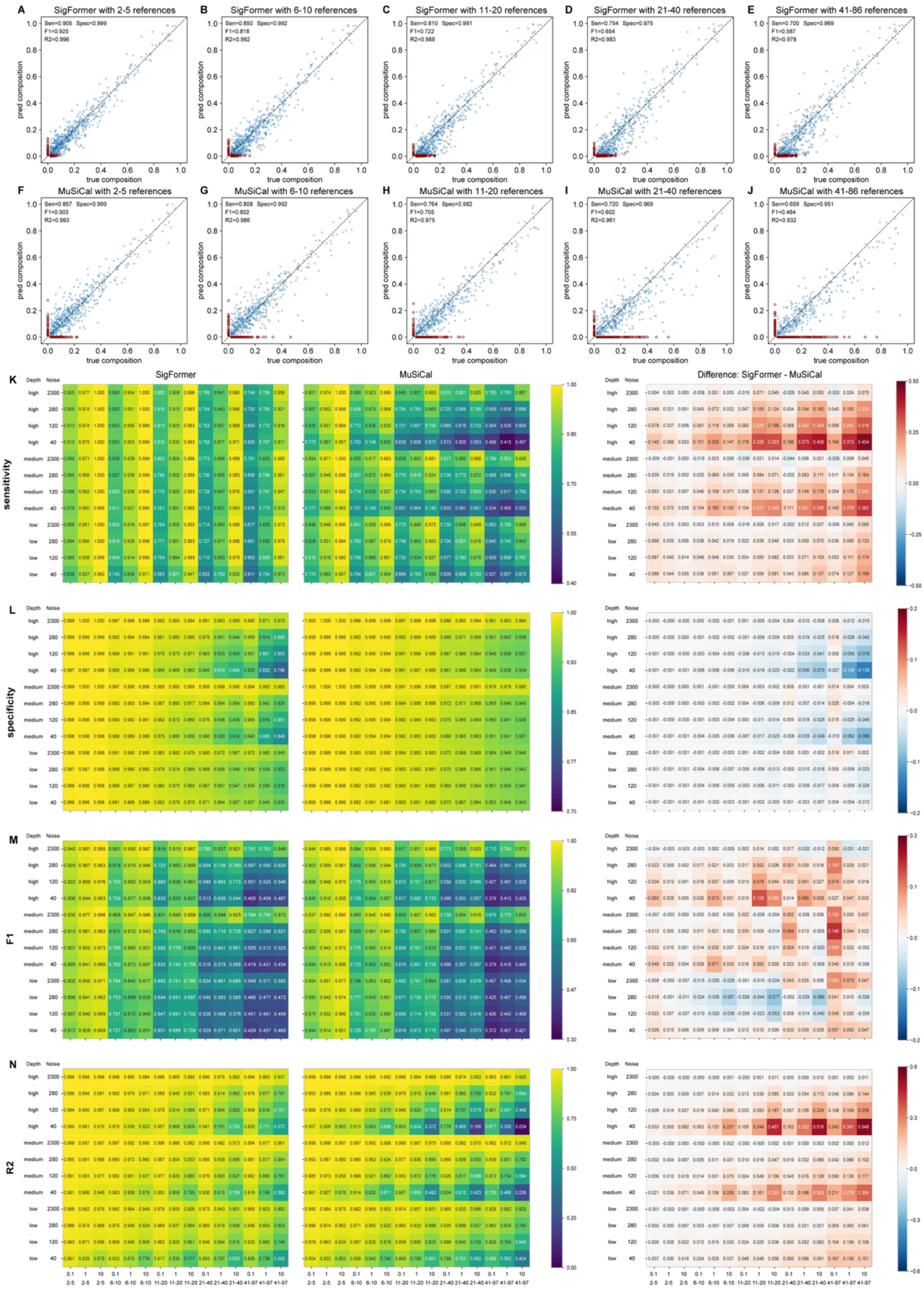

**Supplementary Figure 2: Benchmarking SigFormer and MuSiCal on simulated datasets.** (**A–J**) Representative scatter plots comparing predicted versus ground-truth signature compositions for simulations calibrated to the empirical PCAWG distribution. The example shown corresponds to the setting fitting PCAWG data distribution, specifically: composition Dirichlet alpha = 0.1, medium depth, and Dirichlet–multinomial concentration α = 280 (noise level). Panels are ordered from left to right by increasing reference signature set size, illustrating how performance evolves as the reference pool expands. Each point denotes a (sample, signature) pair; true positives are shown in blue, while false positives and false negatives are highlighted in dark red. (**K–N**) Heatmap summaries across a broader range of simulation conditions. Columns report SigFormer (left), MuSiCal (middle), and their paired difference (SigFormer − MuSiCal; right). Rows correspond to sensitivity, specificity, F1 score, and $R^2$, enabling a systematic comparison of detection accuracy and quantitative agreement across depth, sparsity/composition, noise, and reference-size regimes.

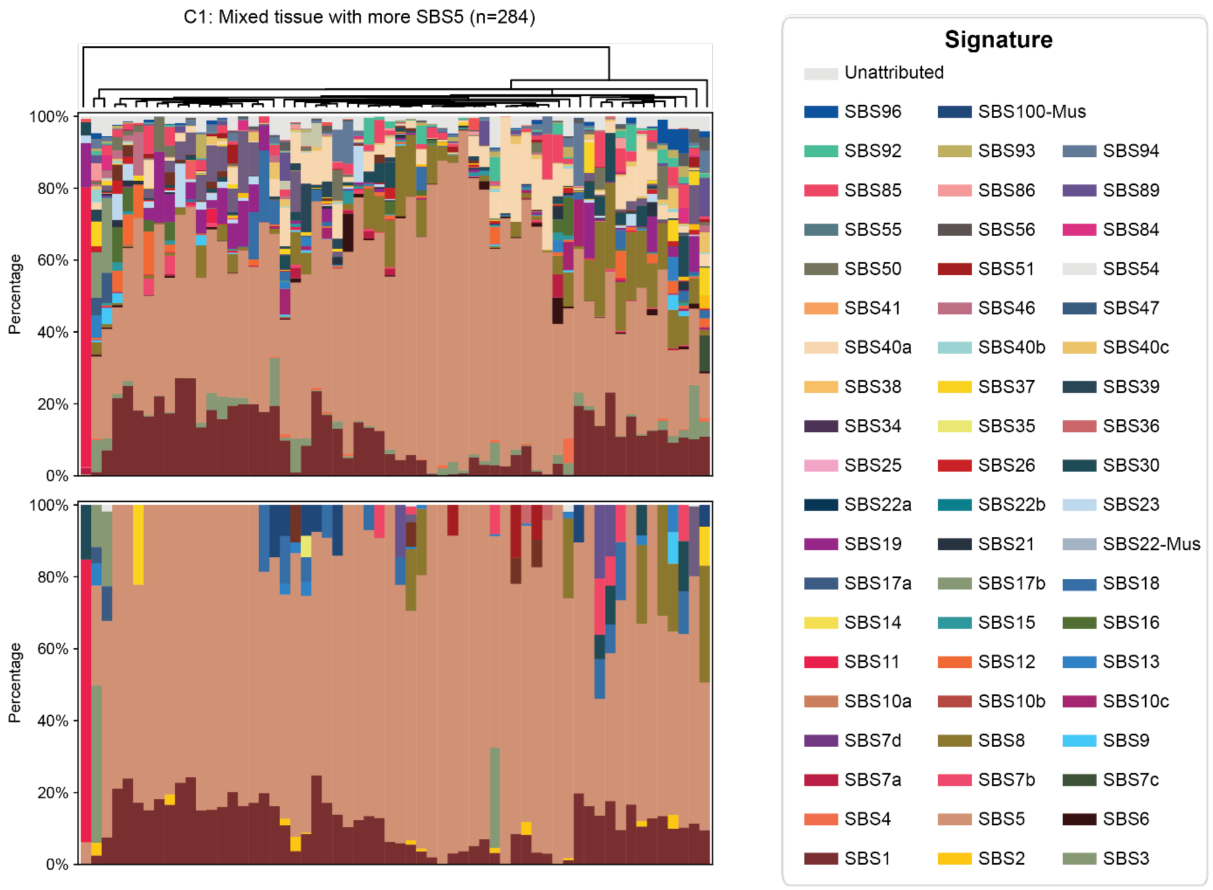


**Supplementary Figure 3:** Signature compositions displayed as stacked barplots for cluster C1, SigFormer (upper) and MuSiCal (lower).

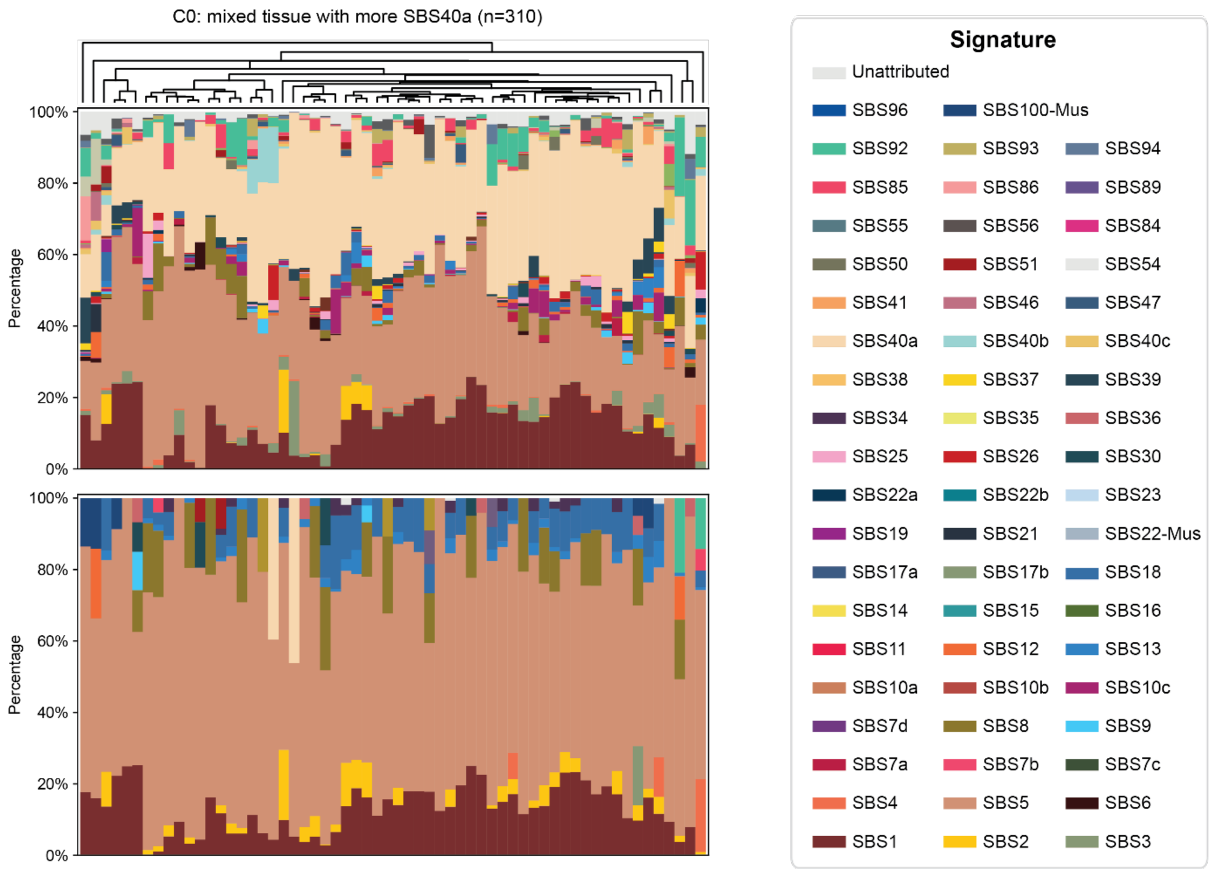


**Supplementary Figure 4:** Signature compositions displayed as stacked barplots for cluster C0, SigFormer (upper) and MuSiCal (lower).

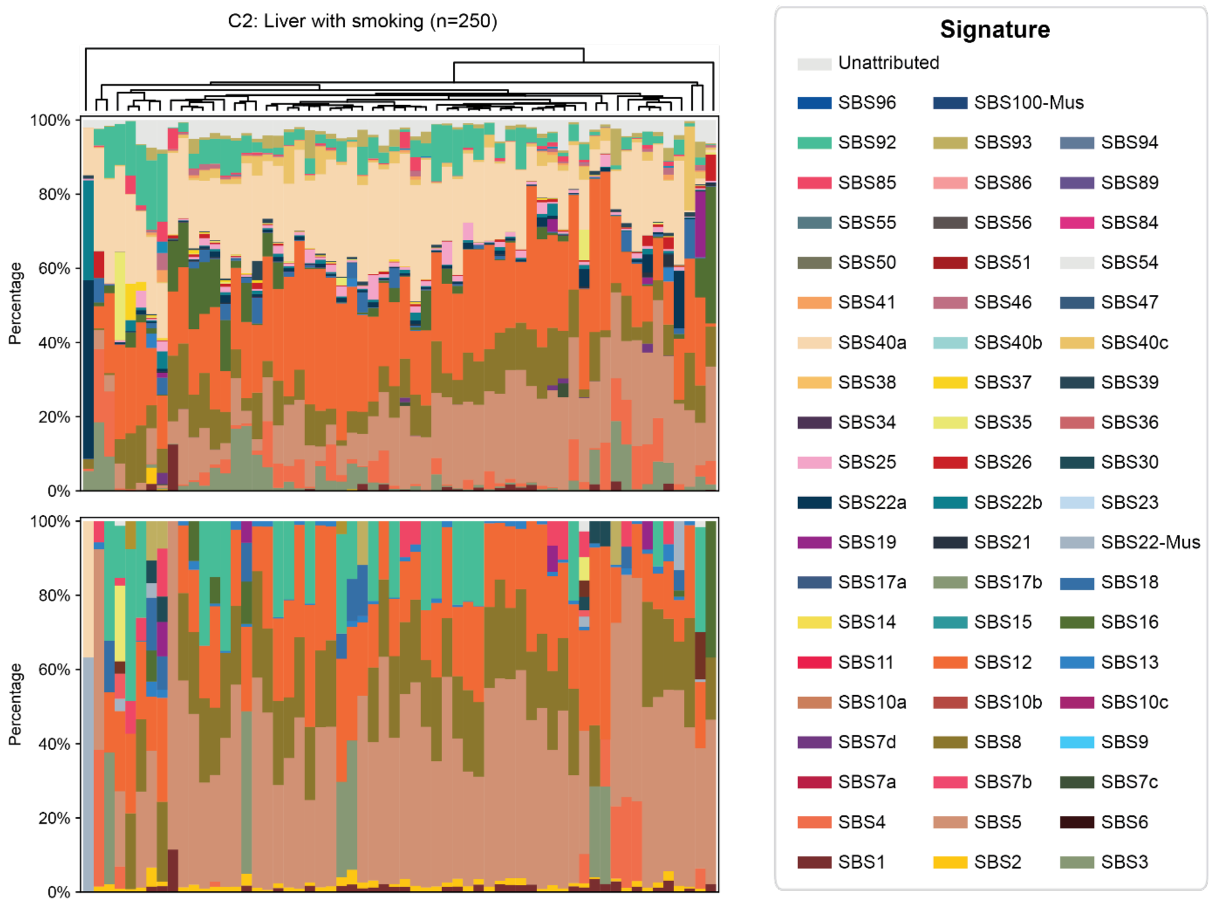


**Supplementary Figure 5:** Signature compositions displayed as stacked barplots for cluster C2, SigFormer (upper) and MuSiCal (lower).

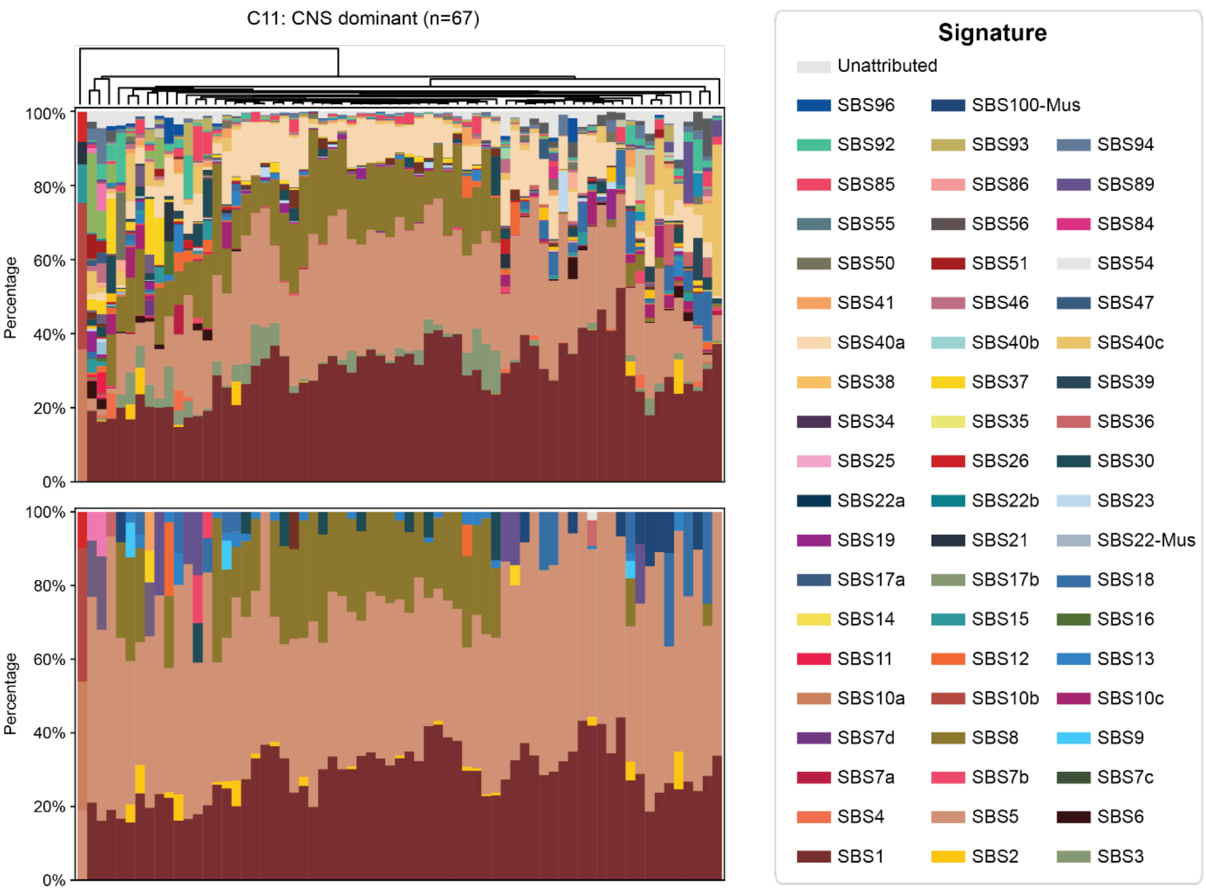


**Supplementary Figure 6:** Signature compositions displayed as stacked barplots for cluster C11, SigFormer (upper) and MuSiCal (lower).

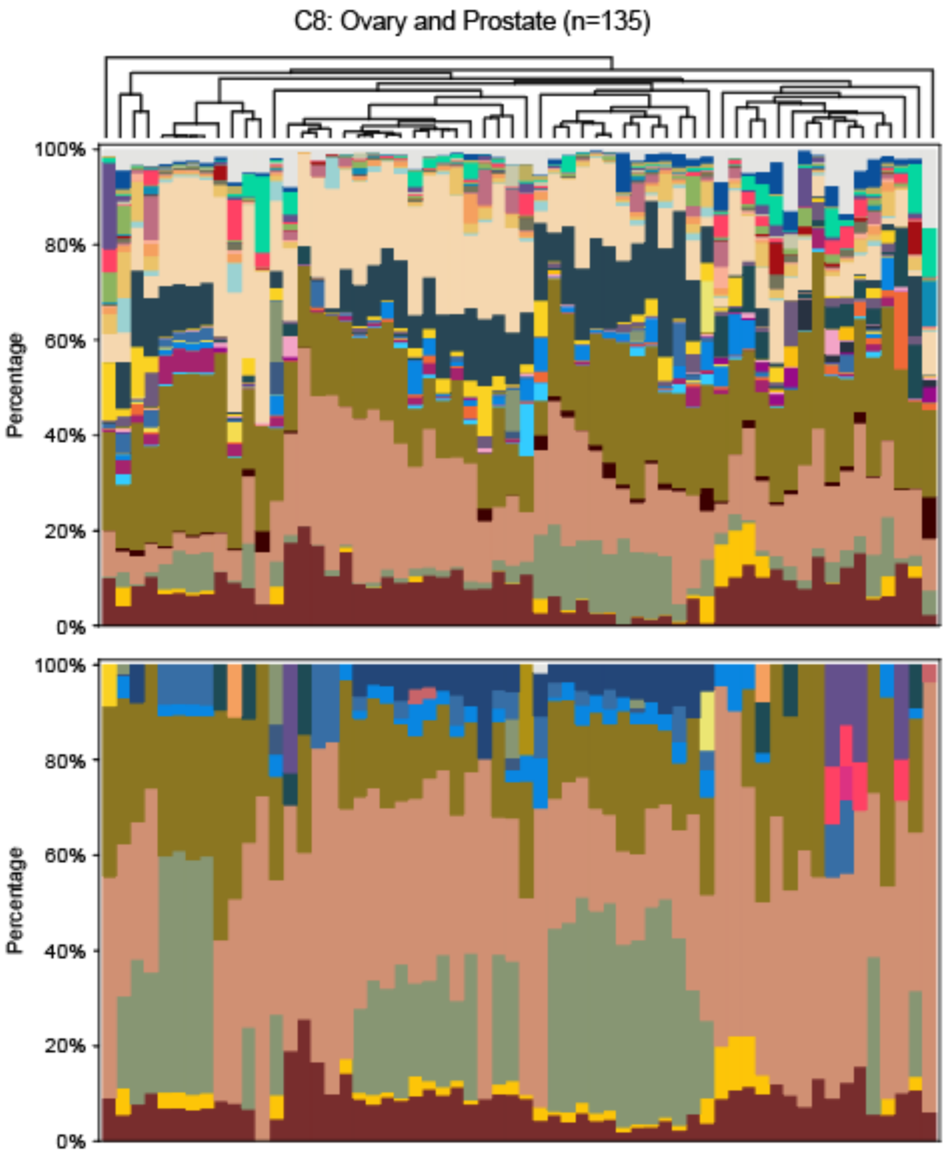


**Supplementary Figure 7:** Signature compositions displayed as stacked barplots for cluster C8, SigFormer (upper) and MuSiCal (lower).

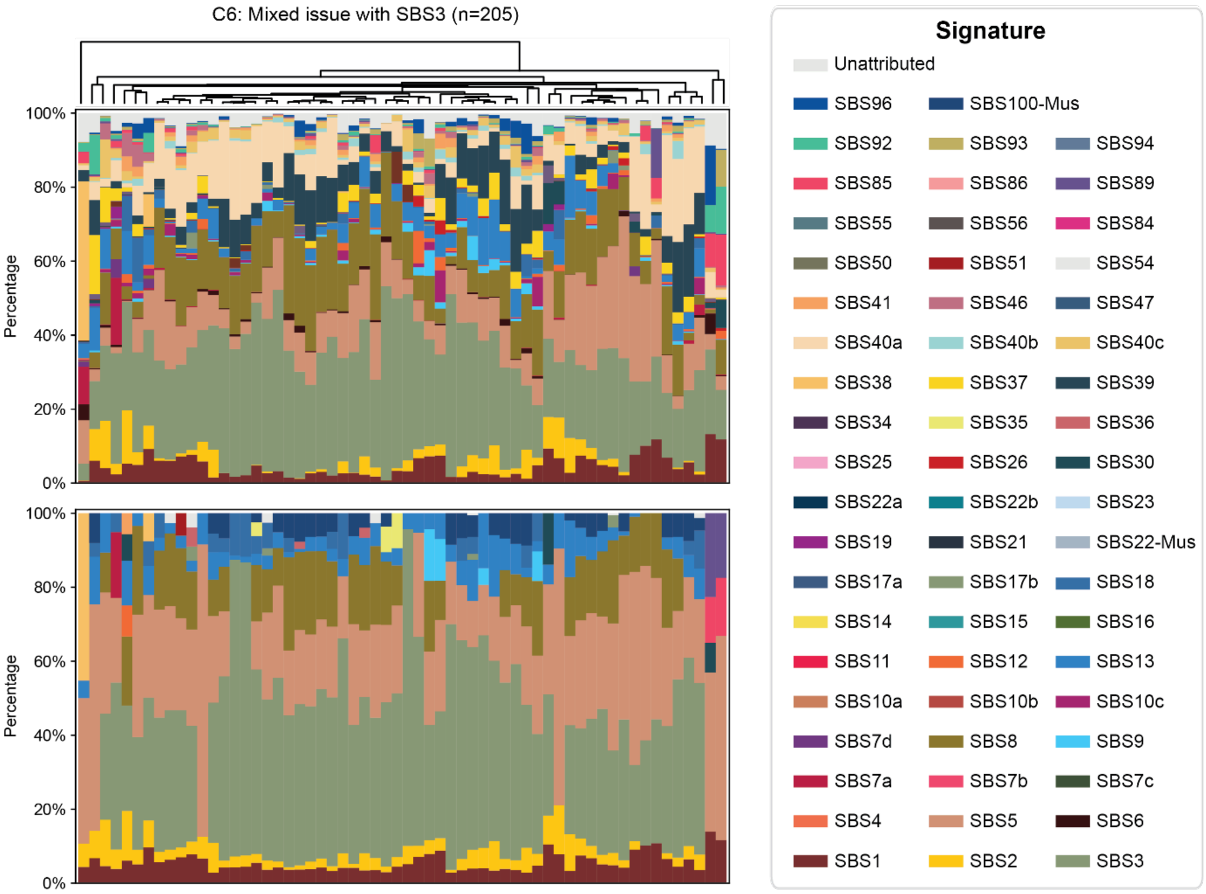


**Supplementary Figure 8:** Signature compositions displayed as stacked barplots for cluster C6, SigFormer (upper) and MuSiCal (lower).

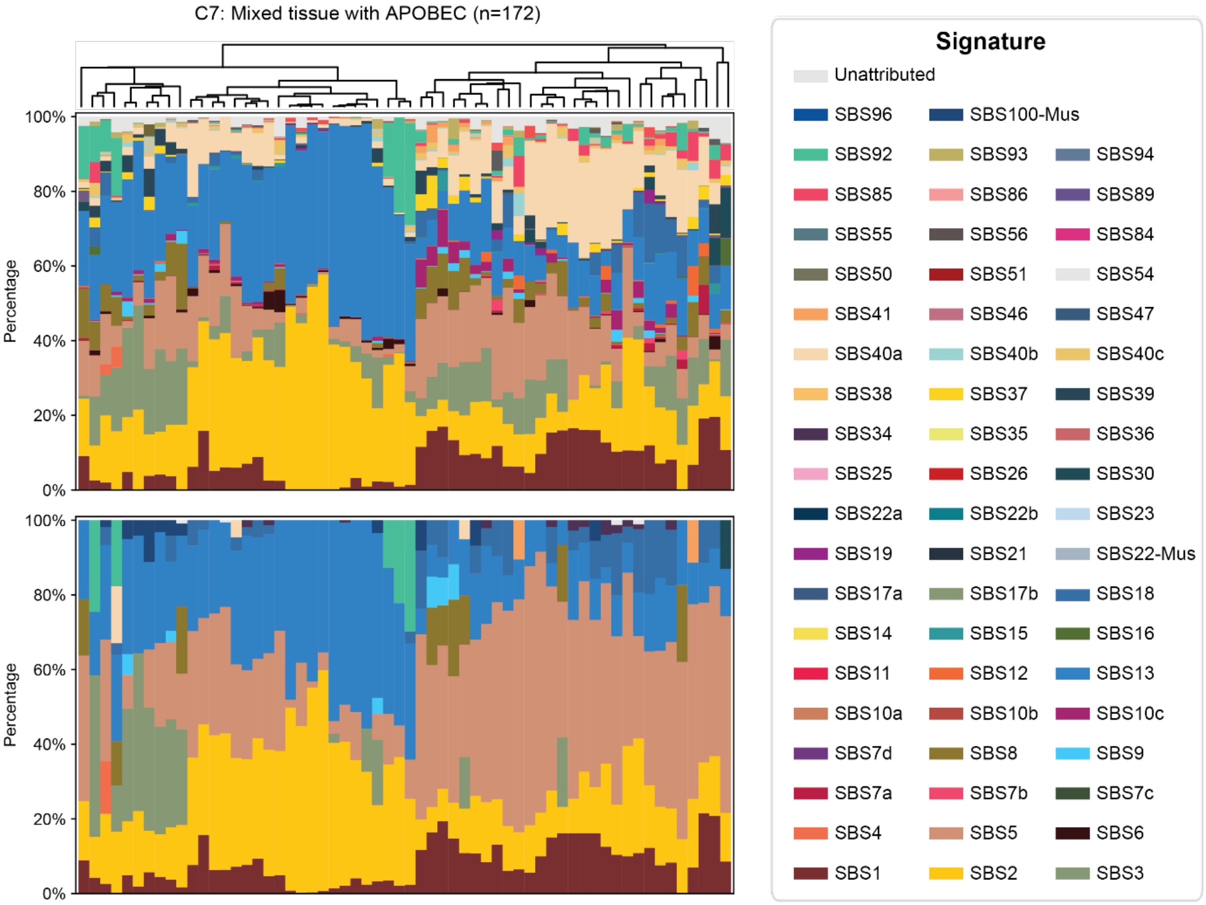


**Supplementary Figure 9:** Signature compositions displayed as stacked barplots for cluster C7, SigFormer (upper) and MuSiCal (lower).

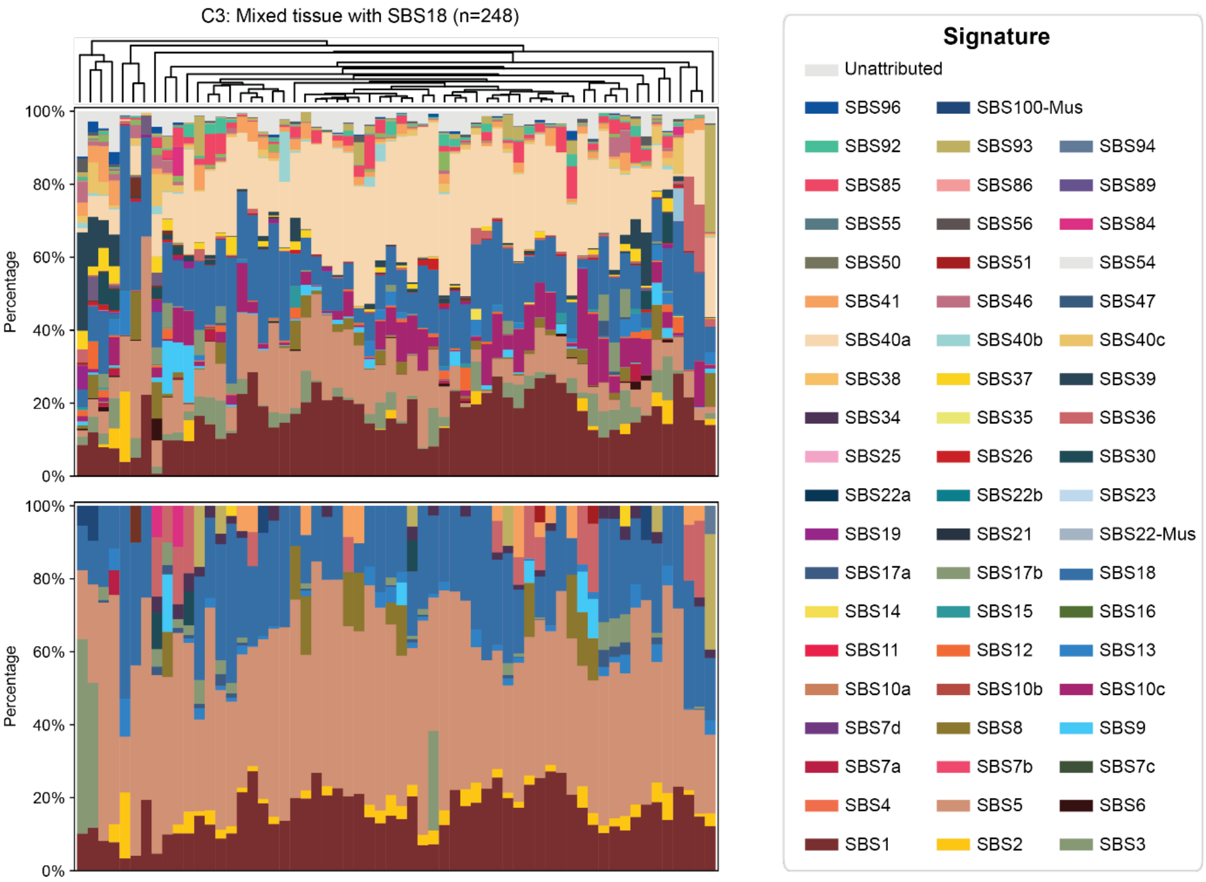


**Supplementary Figure 10:** Signature compositions displayed as stacked barplots for cluster C3, SigFormer (upper) and MuSiCal (lower).

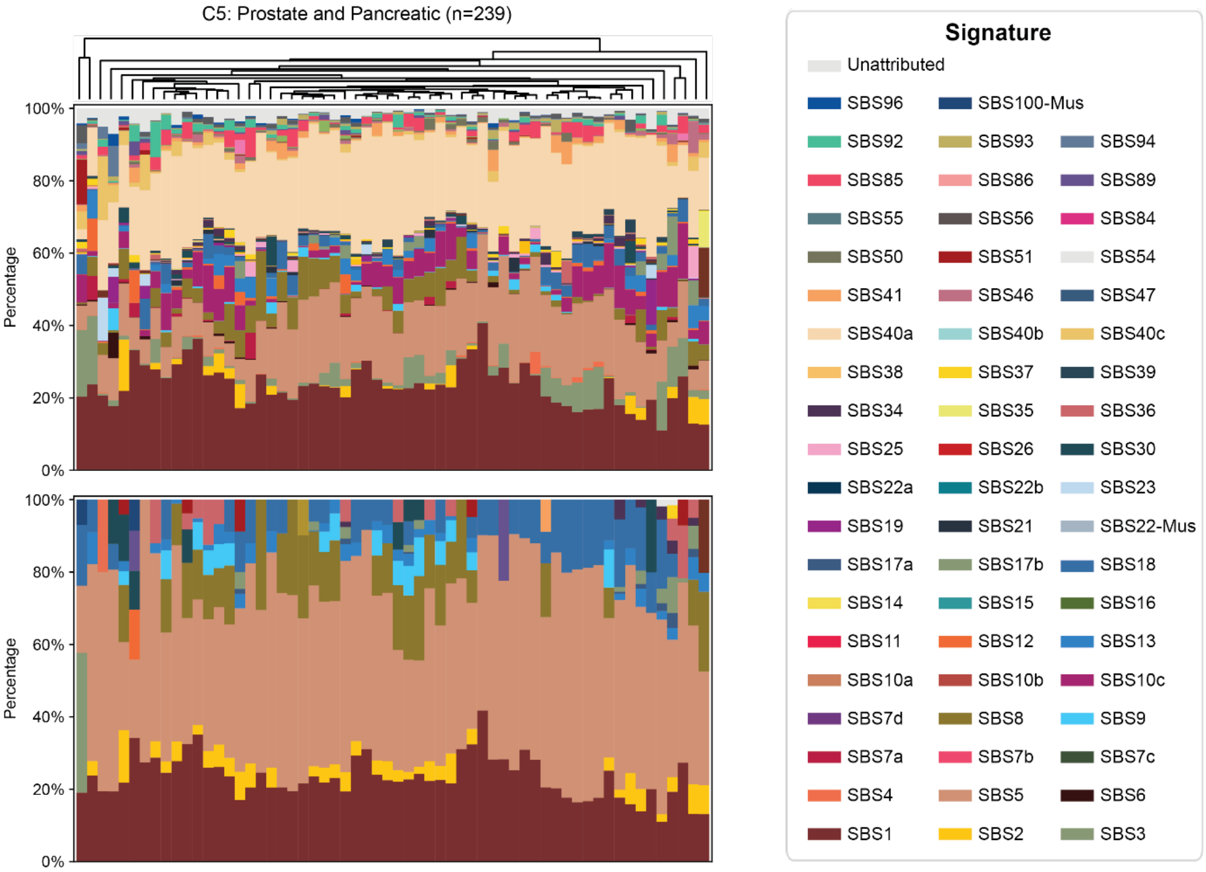


**Supplementary Figure 11:** Signature compositions displayed as stacked barplots for cluster C5, SigFormer (upper) and MuSiCal (lower).

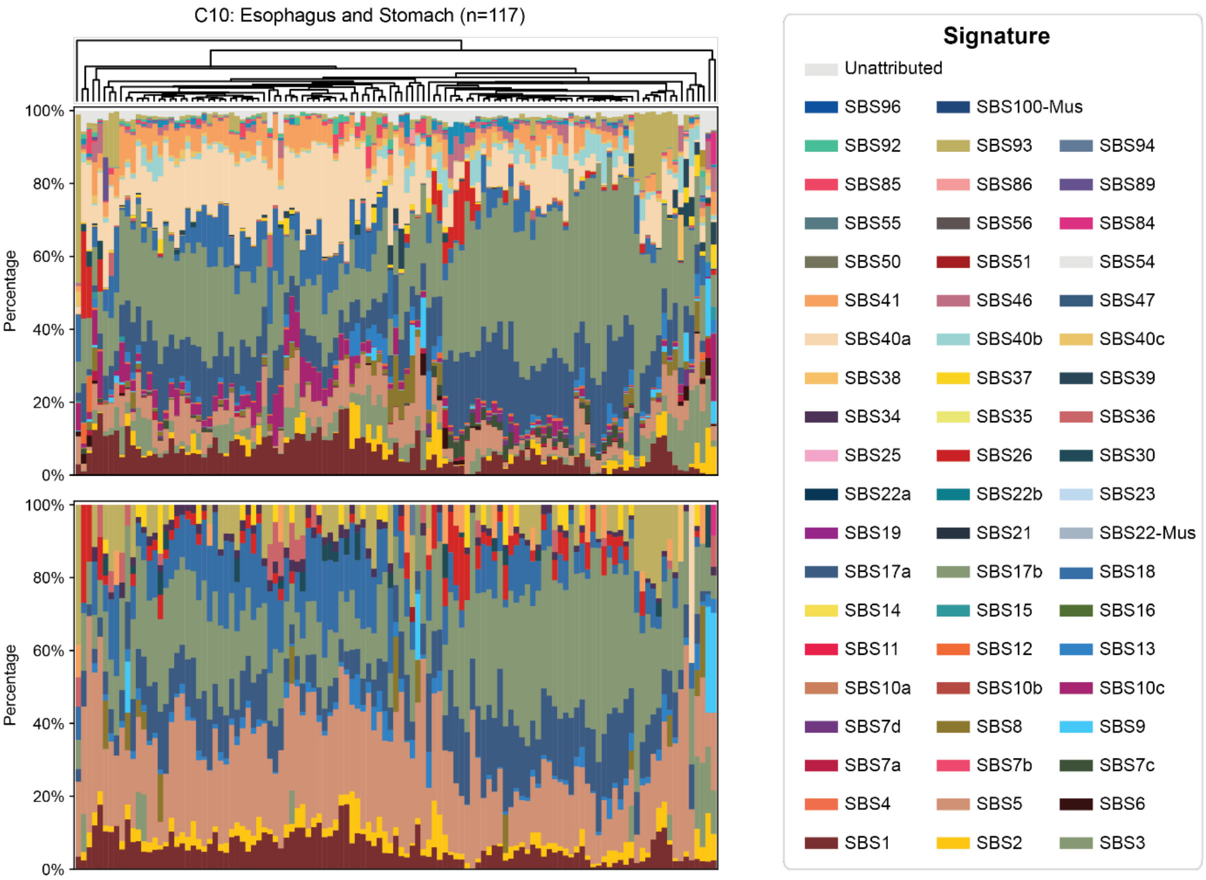


**Supplementary Figure 12:** Signature compositions displayed as stacked barplots for cluster C10, SigFormer (upper) and MuSiCal (lower).

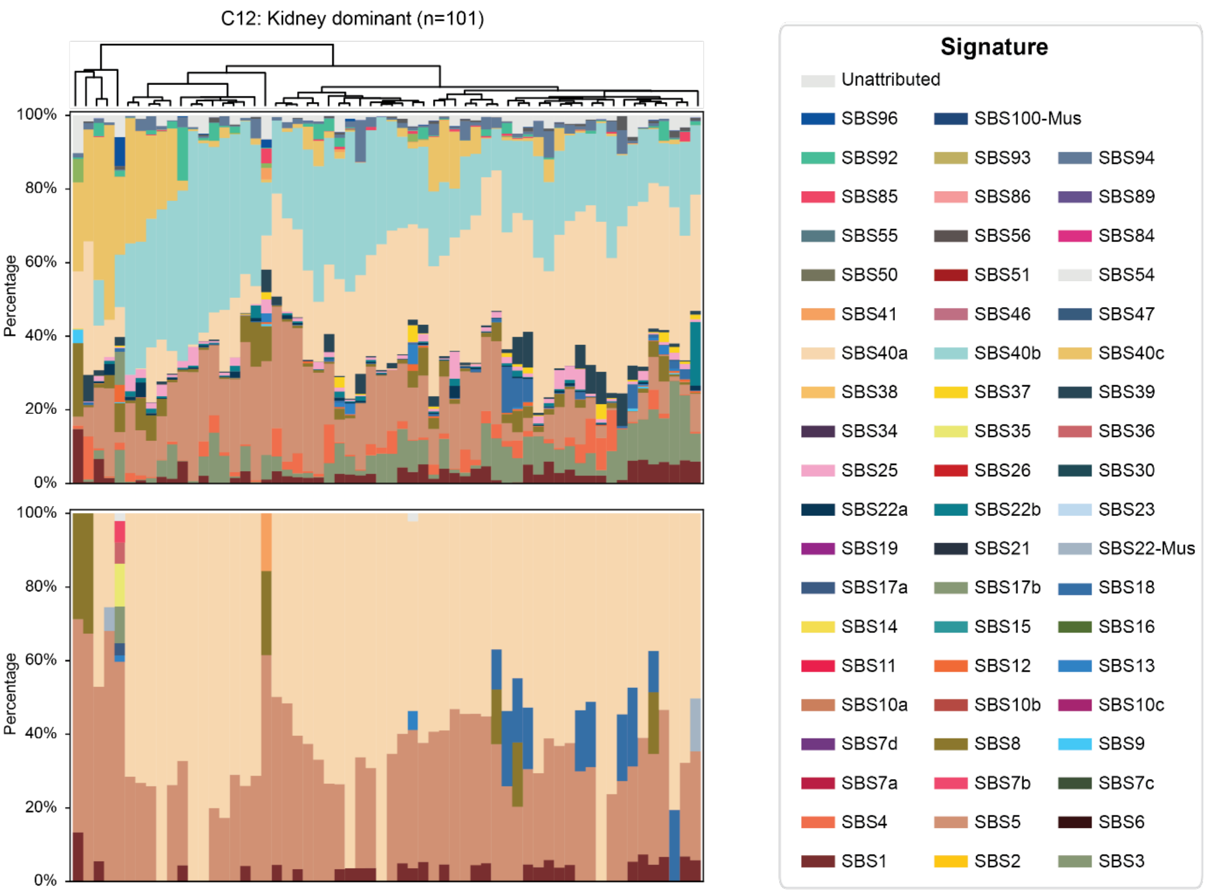


**Supplementary Figure 13:** Signature compositions displayed as stacked barplots for cluster C12, SigFormer (upper) and MuSiCal (lower).

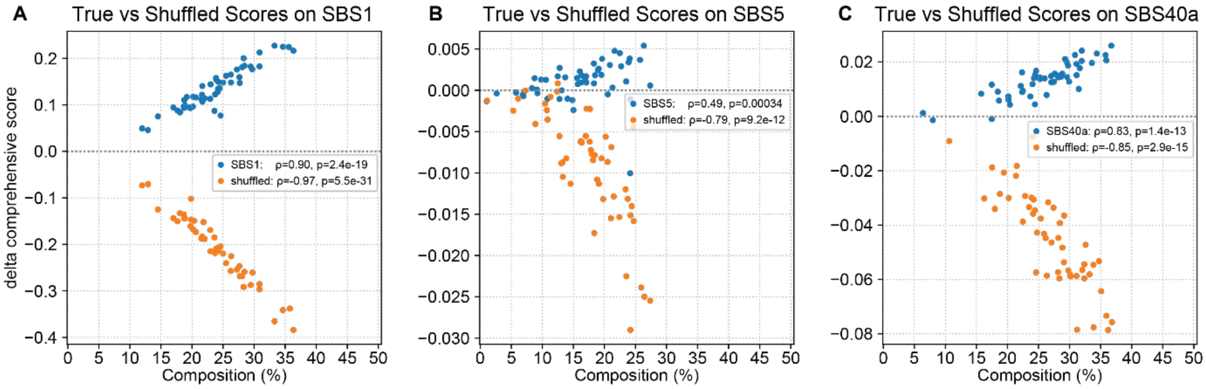


**Supplementary Figure 14:** Evidence-based validation of low-abundance signature detection using a replacement-aware stability test for Cluster C5 (**Supplementary Figure 11**). (**A–C**) Proof-of-concept analysis illustrating the replacement-aware stability test for signatures (SBS1/5/40a). True attributions are compared against replacement controls, where the weights of SBS1/5/40a are redistributed to other reference signatures, followed by re-evaluation of the corresponding signature scores.

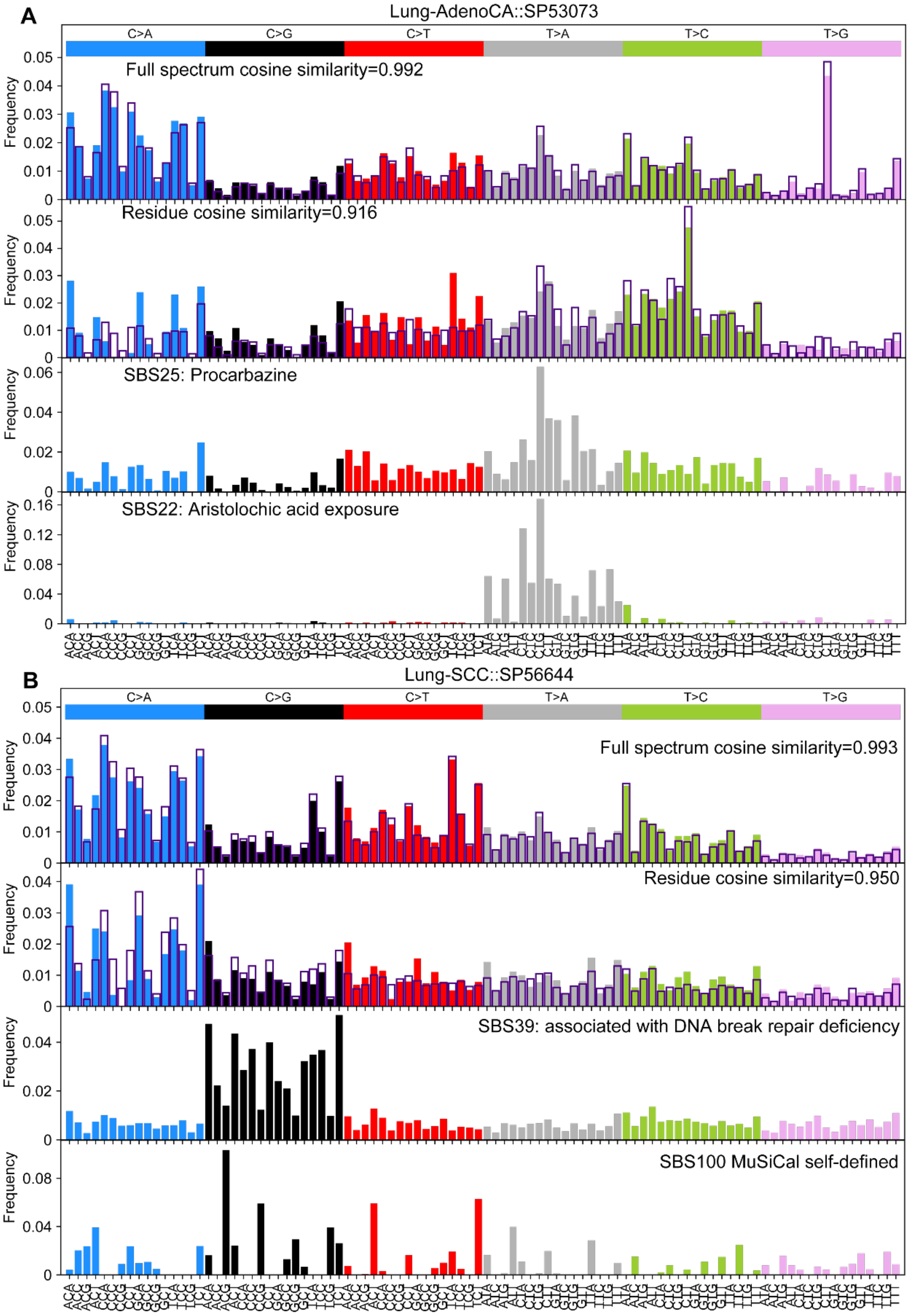


**Supplementary Figure 15:** Spectrum-level inspection for PCAWG Lung samples with smoking signatures (C14). Colored bars show the observed (raw) mutation profil or COSMIC reference profiles, while outlines indicate reconstructions from the inferred signature compositions. Colored bars show the observed (raw) mutation profil or COSMIC reference profiles, while outlines indicate reconstructions from the inferred signature compositions.

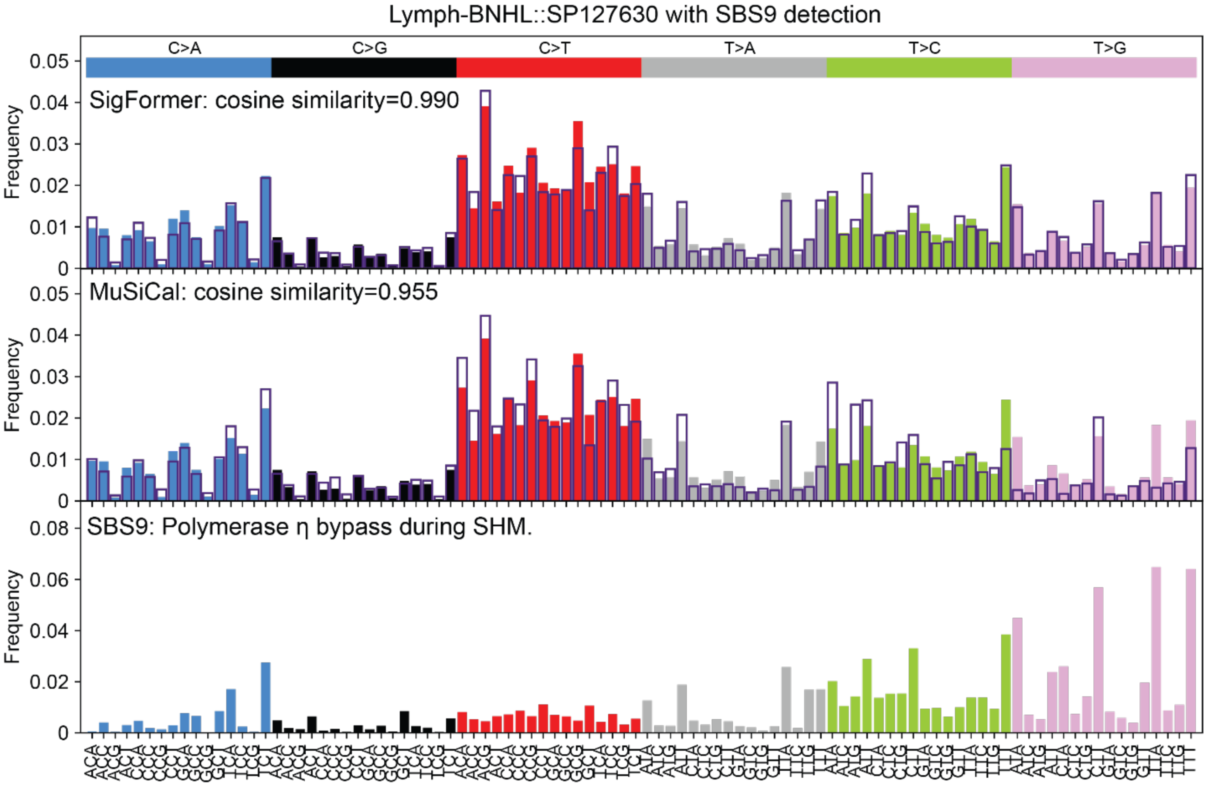


**Supplementary Figure 16:** Spectrum-level inspection for PCAWG lymphocyte cluster (C9). The labels are the same as **Supplementary Figure 15** above.

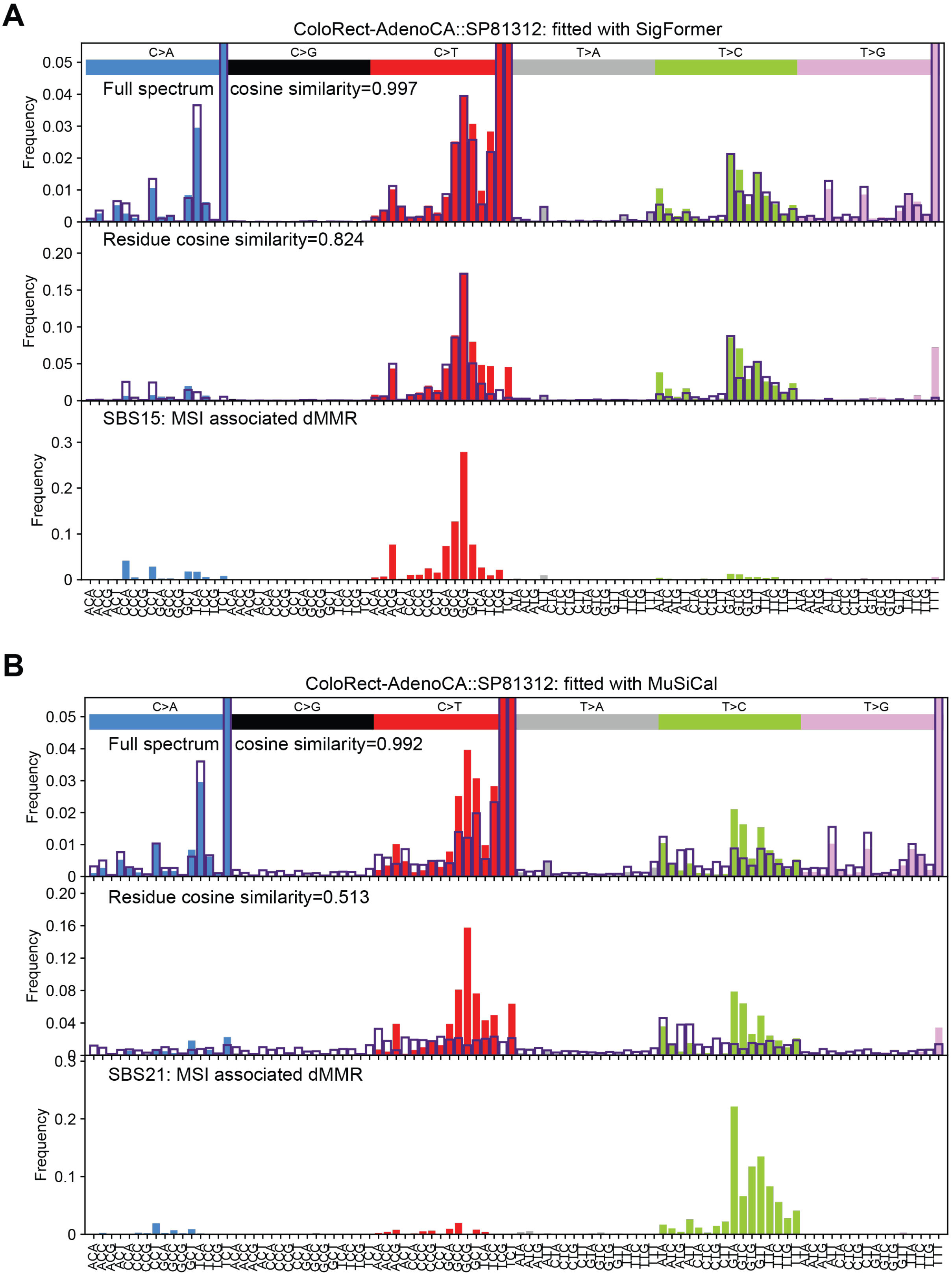


**Supplementary Figure 17:** Spectrum-level inspection for PCAWG cluster C10. The labels are the same as **Supplementary Figure 15** above.

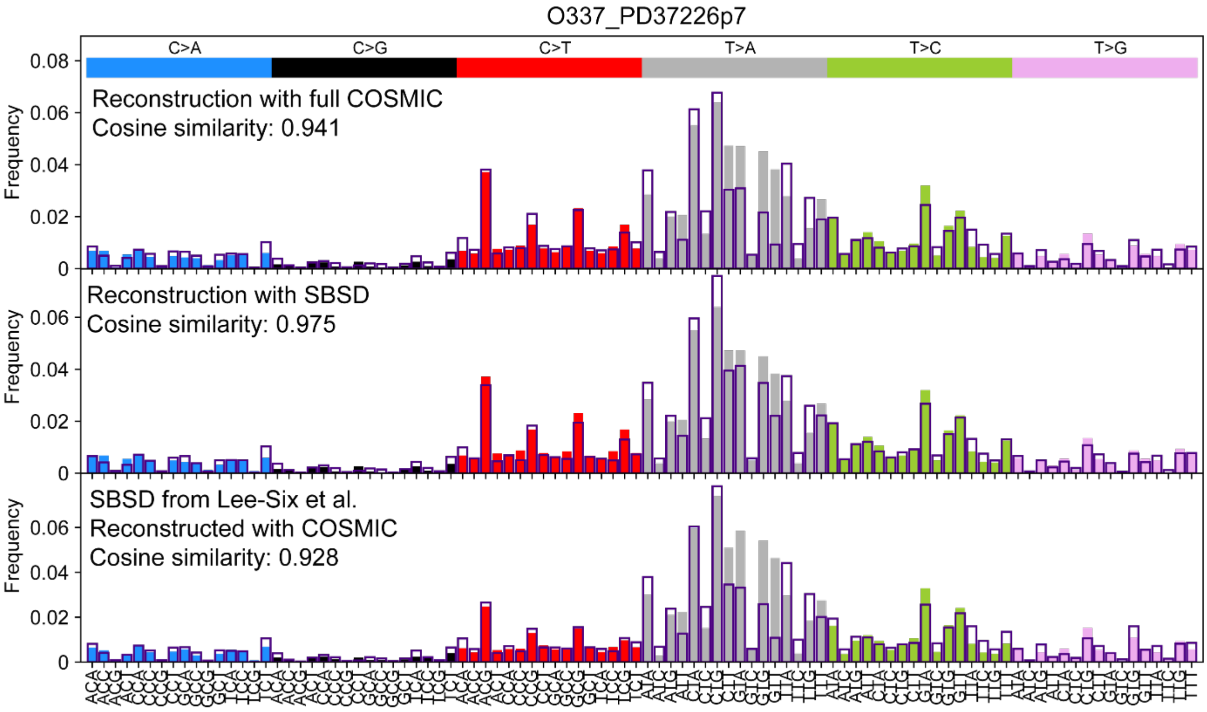


**Supplementary Figure 18:** Spectrum-level inspection for a representative sample receiving multiple chemotherapies from Lee-Six et al. The labels are the same as **Supplementary Figure 15** above.
